# TopoRoot: A method for computing hierarchy and fine-grained traits of maize roots from X-ray CT images

**DOI:** 10.1101/2021.08.24.457522

**Authors:** Dan Zeng, Mao Li, Ni Jiang, Yiwen Ju, Hannah Schreiber, Erin Chambers, David Letscher, Tao Ju, Christopher N. Topp

## Abstract

**Background:** 3D imaging, such as X-ray CT and MRI, has been widely deployed to study plant root structures. Many computational tools exist to extract coarse-grained features from 3D root images, such as total volume, root number and total root length. However, methods that can accurately and efficiently compute fine-grained root traits, such as root number and geometry at each hierarchy level, are still lacking. These traits would allow biologists to gain deeper insights into the root system architecture (RSA).

**Results:** We present TopoRoot, a high-throughput computational method that computes fine-grained architectural traits from 3D X-ray CT images of field-excavated maize root crowns. These traits include the number, length, thickness, angle, tortuosity, and number of children for the roots at each level of the hierarchy. TopoRoot combines state-of-the-art algorithms in computer graphics, such as topological simplification and geometric skeletonization, with customized heuristics for robustly obtaining the branching structure and hierarchical information. TopoRoot is validated on both real and simulated root images, and in both cases it was shown to improve the accuracy of traits over existing methods. We also demonstrate TopoRoot in differentiating a maize root mutant from its wild type segregant using fine-grained traits. TopoRoot runs within a few minutes on a desktop workstation for volumes at the resolution range of 400^3, without need for human intervention.

**Conclusions:** TopoRoot improves the state-of-the-art methods in obtaining more accurate and comprehensive fine-grained traits of maize roots from 3D CT images. The automation and efficiency makes TopoRoot suitable for batch processing on a large number of root images. Our method is thus useful for phenomic studies aimed at finding the genetic basis behind root system architecture and the subsequent development of more productive crops.

## Introduction

Roots are the primary means by which the plant absorbs water and nutrients, and they provide anchorage to the plant. These functions are largely determined by the root system architecture (RSA) [10, 2, 8], which describes both the geometry of individual roots and their hierarchical relationships. Quantifying RSA enables efforts to discover the genetic control of root traits, which can lead to improved crop productivity while minimizing adverse environmental effects [8]. However, RSA is difficult to study owing to roots’ poor accessibility as the “hidden” half of the plant. Traditionally, roots are excavated from the soil, washed, and then measured by hand using devices such as rulers, calipers, and protractors. This process is not only labor-intensive but also prone to human errors. More importantly, many aspects of RSA, particularly those pertaining to lateral roots of higher order, are almost impossible to measure by hand.

Advances in 3D imaging, including X-ray CT, MRI, and optical imaging [20, 27, 15], have allowed root shapes to be captured digitally either after excavation or *in situ*. The availability of such imaging data has paved the way for recent efforts towards computational quantification of root system architecture [23, 19, 28]. However, most image-based root phenotyping methods only compute overall traits such as the volume, depth, convex hull volume, total root length, and root number [33, 6, 14, 5]. Though useful, these traits which are aggregated over the whole root system do not capture the branching structure or the hierarchical organization of individual roots, which provide a much more comprehensive description of RSA. Recently, a system for multi-view scanning and subsequent computational analysis, known as DIRT/3D [12], was proposed to measure detailed traits of maize root crowns. The system can report 18 traits that concern the geometry of the stem (e.g., diameter and depth), the upper whorls (e.g., inter-whorl distance) and their individual nodal roots (e.g., length, angle and diameter). An inherent challenge for multi-view reconstruction, due to occlusion, is resolving densely packed roots. As a result, DIRT/3D does not provide a full root hierarchy of the root crown beyond the nodal roots.

To our knowledge, DynamicRoots [32] is the only published and validated root phenotyping method that produces a full branching hierarchy and root traits associated with each hierarchy level in 3-dimensions. DynamicRoots is designed for a time-series of root systems grown in transparent gel [17]. These seedling-stage root systems tend to have a relatively simple geometry and structure, which makes it possible to obtain high quality 3D reconstructions using multi-view imaging [39]. DynamicRoots first employs graph analysis on the voxelized root system at each time-point to identify the root branches. Hierarchical relations among the branches are first determined by the length of the branches and then refined by the time function obtained by aligning root architectures across time. However, DynamicRoots makes two assumptions that limit its application to other types of root images. First, the input shapes can only have a mild amount of topological errors, including disconnected root components (e.g. due to limits in resolution), root branches forming loops due to touchings (e.g. due to limits in resolution and noise), and voids of space surrounded by the surface of the root (e.g. due to high optical density of the root). Second, the time series must be dense enough to observe the correct root hierarchy. Neither assumption may hold for other imaging setups, such as X-ray CT or MRI scans of complex roots, which may lead to numerous topological errors after image segmentation and have few, and often a single, time point available.

In this work, we present TopoRoot, a method for obtaining the complete root hierarchy and computing the associated traits from a single 3D X-ray CT scan of excavated maize root crowns. TopoRoot adopts a multi-step approach to treat the topological errors in the input image, and it computes the hierarchy and traits using a high-fidelity skeleton representation of the root system architecture. Our method builds on state-of-the-art algorithms from computer graphics and introduces customized heuristics tailored to the image features and the maize root structure.

On a set of X-ray CT scans of 45 excavated maize root crowns where manual measurements of nodal root counts were made, TopoRoot shows dramatic improvements in accuracy compared to DynamicRoots. As a demonstration of the utility of TopoRoot, several of TopoRoot’s fine-grained traits showed an ability to differentiate between two groups of genetically differentiated species within this data set. On another set of 495 maize root images simulated by OpenSimRoot [24] with varying age, complexity, and noise level, TopoRoot exhibits improved accuracy in a variety of coarse-grained and fine-grained traits over GiaRoots [33] and DynamicRoots.

TopoRoot is completely automated and has only a few parameters to set. On a standard desktop computer, TopoRoot runs within a few minutes for images at the resolution range of 400^3. This makes TopoRoot suited for batch processing a large set of images in a high-throughput analysis pipeline. The software package is freely distributed on GitHub with our test dataset.

## Material and Methods

### Data preparation

The main data set used for validation consists of 59 maize roots which were planted in June 2020 at Planthaven Farms in O’Fallon, Missouri (latitude 38.84871204483824, longitude - 90.68711352048403) in silt loam soil. Fields received fertilization with ammonium nitrate. Seeds were planted using jabtype planters into 3.65-m long rows (~25.4-cm within row spacing) on 0.9144-meter row-to-row spacing. The maize plants (Rt1-2.4 MUT) that had mutation on the *Rootless1* gene are known to have decreased nodal root counts [16] compared to their wild-type counterparts (Rt1-2.4 WT). After 54-57 days of growth, the roots were excavated using the Shovelomics protocol [34] in September 2020 and washed to remove soil. We used an X5000 X-ray imaging system and efX-DR software (NSI, Rogers, MN) to collect X-ray computed tomography (XRT) data (pictured in Figure 1). The X-ray source was set to a voltage of 70kV, current of 1700μA, and focal spot length of 119μm. Samples were clamped and placed on a turntable for imaging at a magnification of 1.17X and 10 frames per second (fps), collecting 1800 16-bit digital radiographs over a 3 minute scan time. efX-CT software was used to reconstruct the scan into a 3D volume at 109μm voxel resolution. This volume was exported as a 16-bit RAW volume. Following imaging, manual counts were collected for the nodal roots. Each sample was dissected starting at the highest node (stalk end) moving downward to the root tips. Only attached roots were counted towards the total number of developed roots at each node. Finally, 14 scans were removed from the analysis due to excessive soil present in the imaging and missing whorls. The remaining 45 scans and their manual nodal root counts provide validation for TopoRoot’s computed values.

**Fig. 1.**
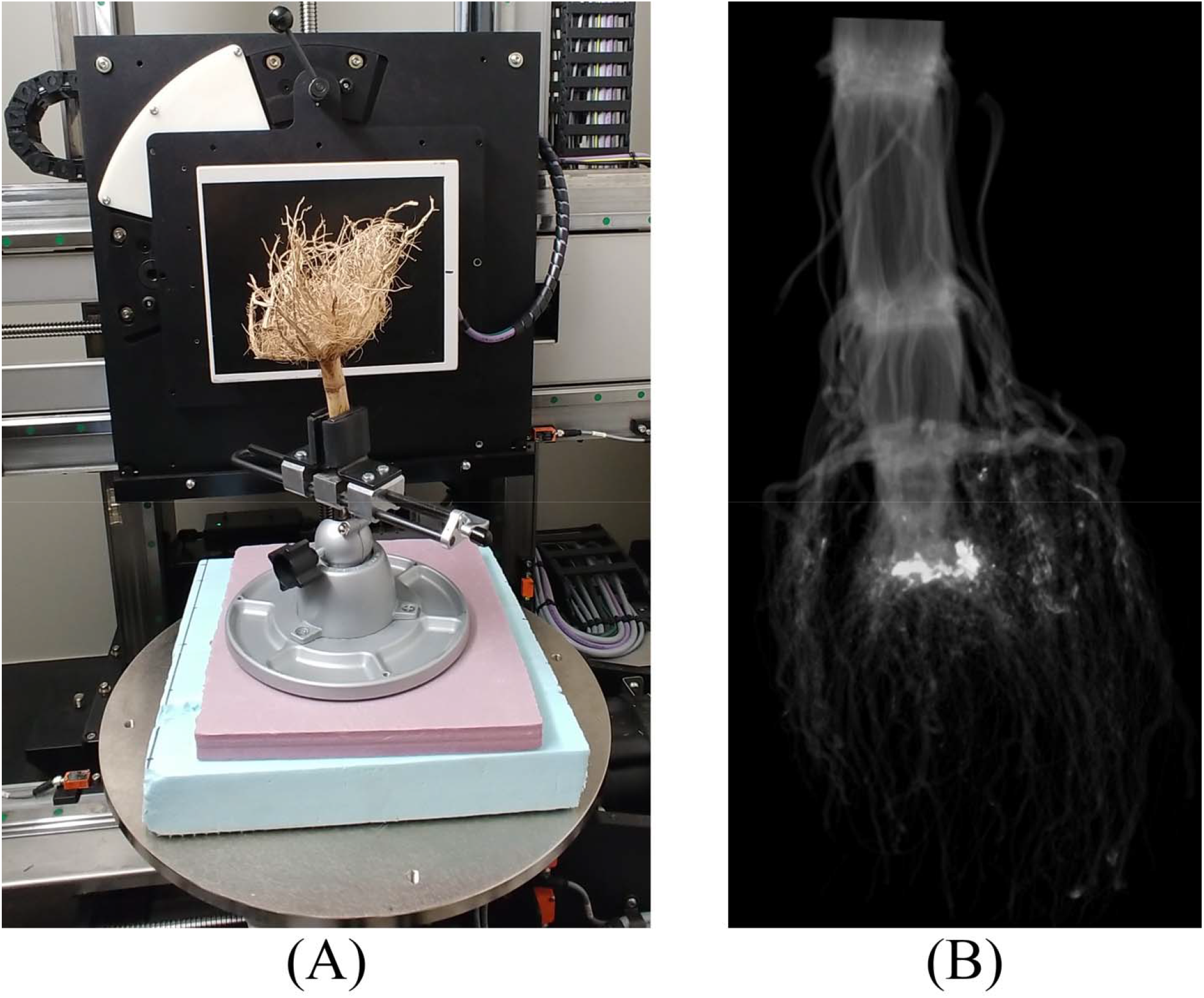
Picture of the X-ray setup. (A) Each maize root crown was clamped and placed on a turntable, which was rotated for 3 minutes while radiographs were collected at a rate of 10 frames per second. (B) efX-CT software was used to reconstruct the scan into a 3D grayscale volume (pictured).

It is generally difficult to obtain manual measurement of fine-grained root traits beyond counting the nodal roots. To validate other fine-grained traits produced by our method, we supplement the real data set with a systematically created set of simulated root images. We adopt OpenSimRoot [24], a highly customizable 3D root growth simulation software that has been widely used in modeling and visualizing root growth [11, 31, 7]. We used OpenSimRoot to create 55 maize roots ranging in days of growth from 30 to 40 days, numbers of nodal roots ranging from 31 to 69, number of whorls from 5 to 6, and lateral root branching frequency from 0.3 to 0.7 cm / branch. The diameter of the stem was set to be 2 cm, starting diameter for nodal roots is 0.3 cm (gradually decreasing to 0.1 cm after 10 days of growth), lateral roots is 0.04 cm, and fine lateral roots is 0.02 cm. OpenSimRoot provides a detailed hierarchy for each of the simulated roots, from which we obtain the ground-truth traits.

For each simulated root, we create a 512^3 image by computing the signed distance field from the surface of the root using the method of [35] with the inside of the surface having positive values and the outside having negative values. To simulate various levels of image noise, we randomly perturb the distance value at each voxel, with the amount of perturbation ranging from 0 to 0.08 cm in 0.01 increments. This results in 9 images at increasing noise levels for each of the 55 roots, and thus 495 images in total. Figure 2 shows images of one simulated root (at day 40) at different noise levels. Note that the amount of geometric irregularity and topological noise (e.g., disconnected components and loops) increase with the noise level.

**Fig. 2.**
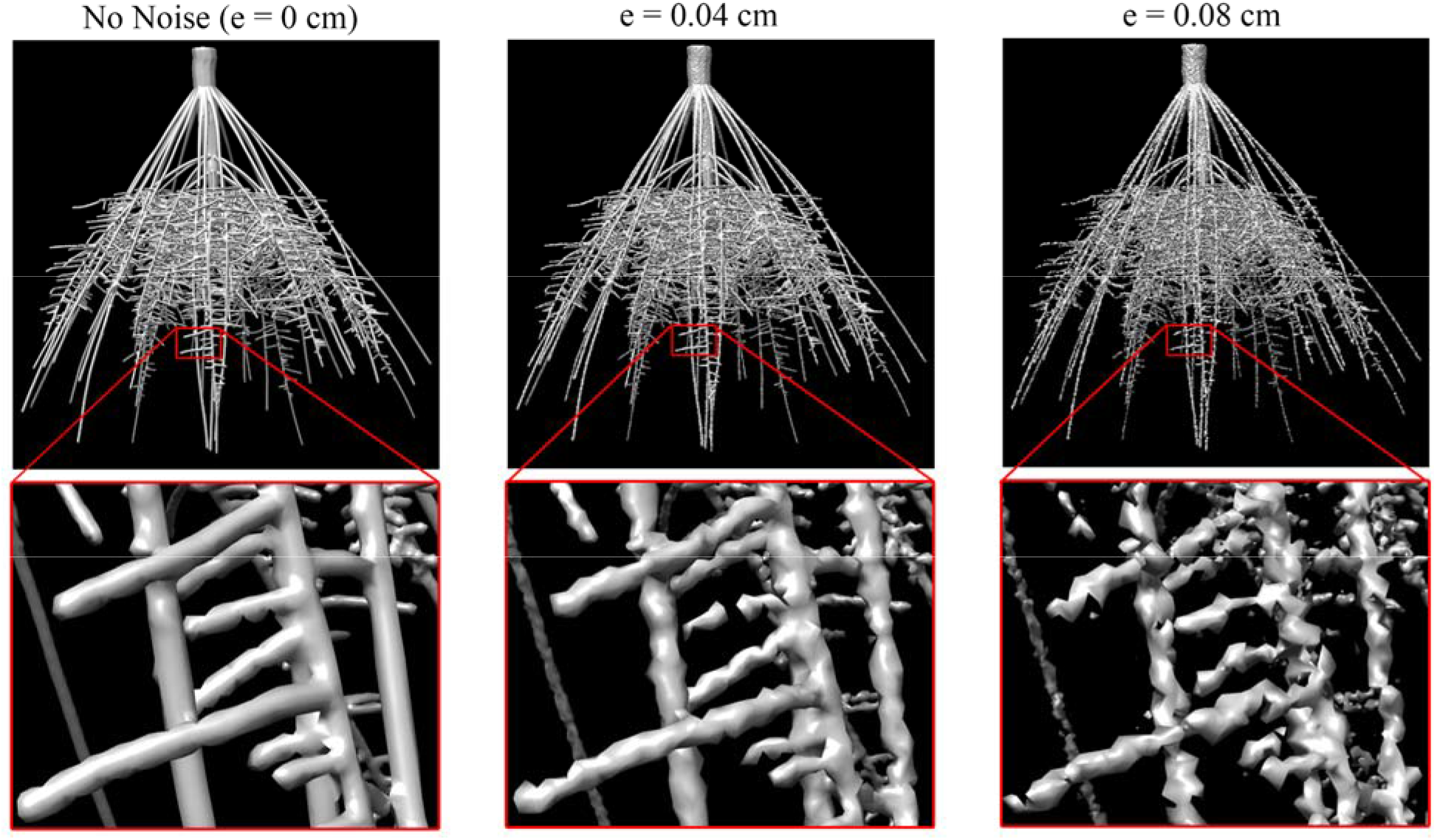
Iso-surfaces (at level 0) of simulated maize root images with increasing amounts of noise (e).The example is at 40 days of growth. The closeups show fine lateral roots. With increased noise, the roots exhibit less regular geometry and more topological errors (e.g., disconnections and loops).

### Thresholding

Although the X-ray images have good contrast between roots and air, due to the spatial variation in the contrast, limited resolution and noise, there is typically no common threshold that can accurately capture both thicker (e.g., nodal) and thinner (e.g., lateral) roots. To obtain the best result, our method therefore asks the user to provide three thresholds, visualized in Figure 3. The shape threshold provides the best balance between capturing thick and thin roots, but it may contain many topological errors (e.g., disconnected thin roots and merged thick roots). We additionally ask for a kernel threshold (the lowest value that avoids merging of thick roots) and a neighborhood threshold (the highest value that avoids disconnecting thin roots), such that the increasing ordering of the three thresholds are neighborhood, shape, and kernel. Our method will attempt to fix topological errors at the shape threshold guided by the neighborhood and kernel thresholds. For this data set, we set the shape threshold to be 0, kernel threshold to be 0.03 and neighborhood threshold to be −0.15.

**Fig. 3.**
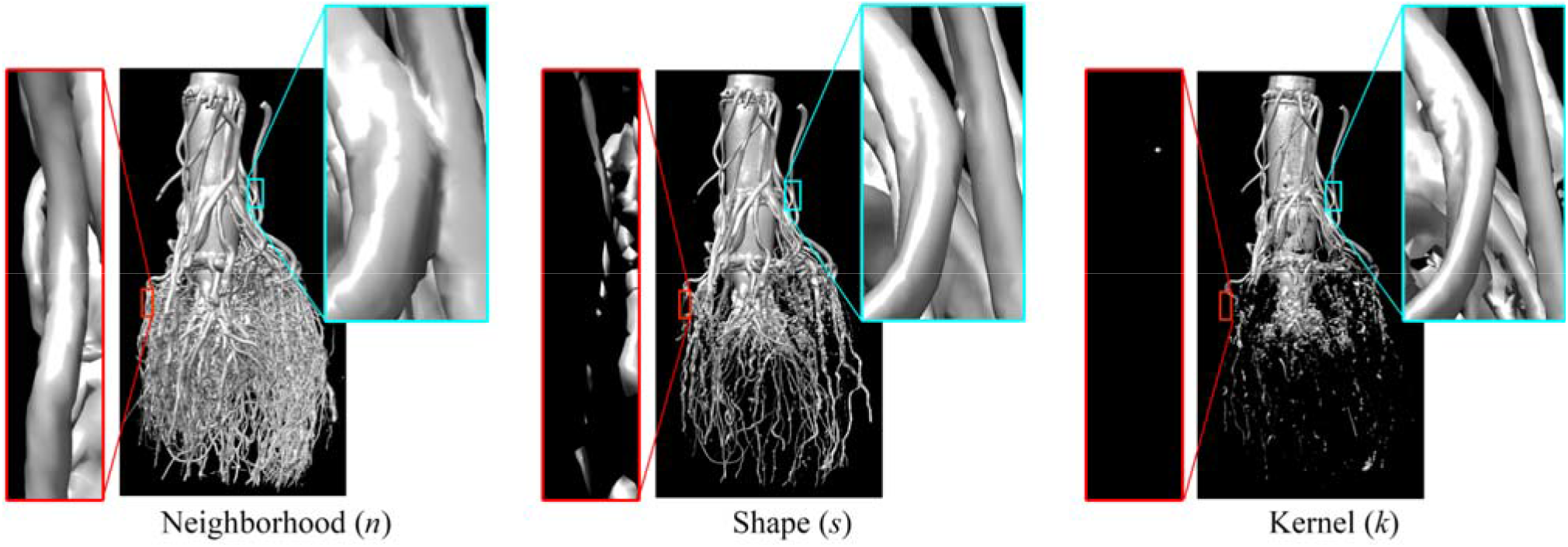
Iso-surfaces *n*, *s*, and *k*, as visualized for an X-ray image of a maize root. The red closeup shows a fine lateral root which is connected in *n*, but disconnected in *s* and which has disappeared in *k*. The light blue closeup shows two nodal roots which are merged together in *n* and *s*, but separated in *k*.

### Overview

Our pipeline takes as input a 3D grayscale image with three thresholds (shape, kernel and neighborhood) and produces a hierarchy and associated traits in four steps (Figure 4):

1. Topological simplification: Most topological errors on the iso-surface at the shape threshold are removed by combining a global optimization algorithm [38] and a heuristic that fills in the hollow space inside the stem and other thick roots.
2. Skeletonization: A geometric skeleton capturing the branching structure is computed by first running an existing skeletonization algorithm [36, 37] and then removing cycles on the skeleton using a minimum-spanning tree.
3. Inferring hierarchy: Each skeleton branch is associated with a hierarchy level using a heuristic that favors a shallow hierarchy while encouraging longer branches at lower levels.
4. Computing traits: A suite of root traits, such as the count, lengths, angles, thickness, and tortuosity, are computed from the skeleton at each level of the hierarchy.

These steps are detailed in the next sections.

**Fig. 4.**
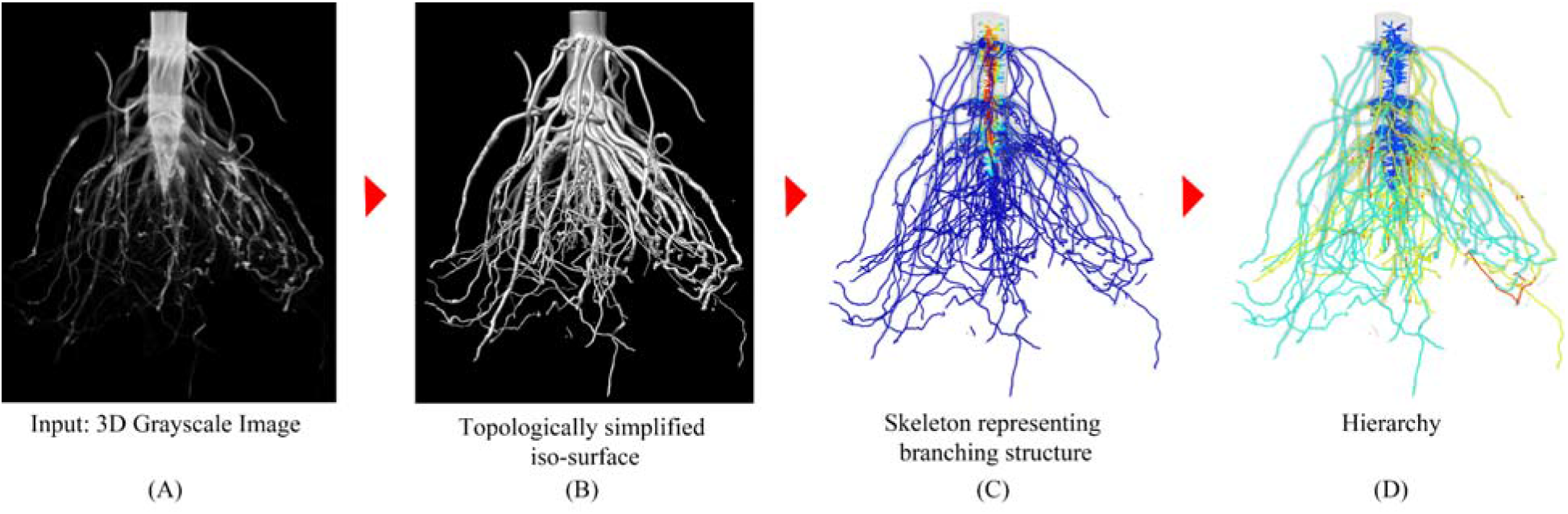
The pipeline of TopoRoot for computing fine-grained traits from a 3D X-ray image. Beginning from a 3D grayscale image (A), TopoRoot first simplifies the topological complexity of the iso-surface (B), then it creates a geometric skeleton capturing the branching structure (C), from which a hierarchy is obtained (D) and the traits are subsequently computed.

### Topological simplification

As shown in Figures 1 and 2, iso-surfaces of input images often contain numerous topological errors, including disconnections, loops, and voids. These errors make it challenging to infer the branching structure of the root system. We use a recent algorithm [38] to maximally remove topological errors. Given the iso-surfaces at three thresholds (shape, kernel and neighborhood), the algorithm attempts to remove all topological features on the iso-surface at the shape threshold that are not present on either of the other two iso-surfaces. The algorithm allows both addition and removal of contents to the shape’s iso-surface, and it tries to minimize these geometric changes to achieve topological simplification. This algorithm can effectively connect broken branches (if they are contiguous at the neighborhood threshold) and split merged branches (if they are separate at the kernel threshold), as shown in Figure 5A and 5B.

**Fig. 5.**
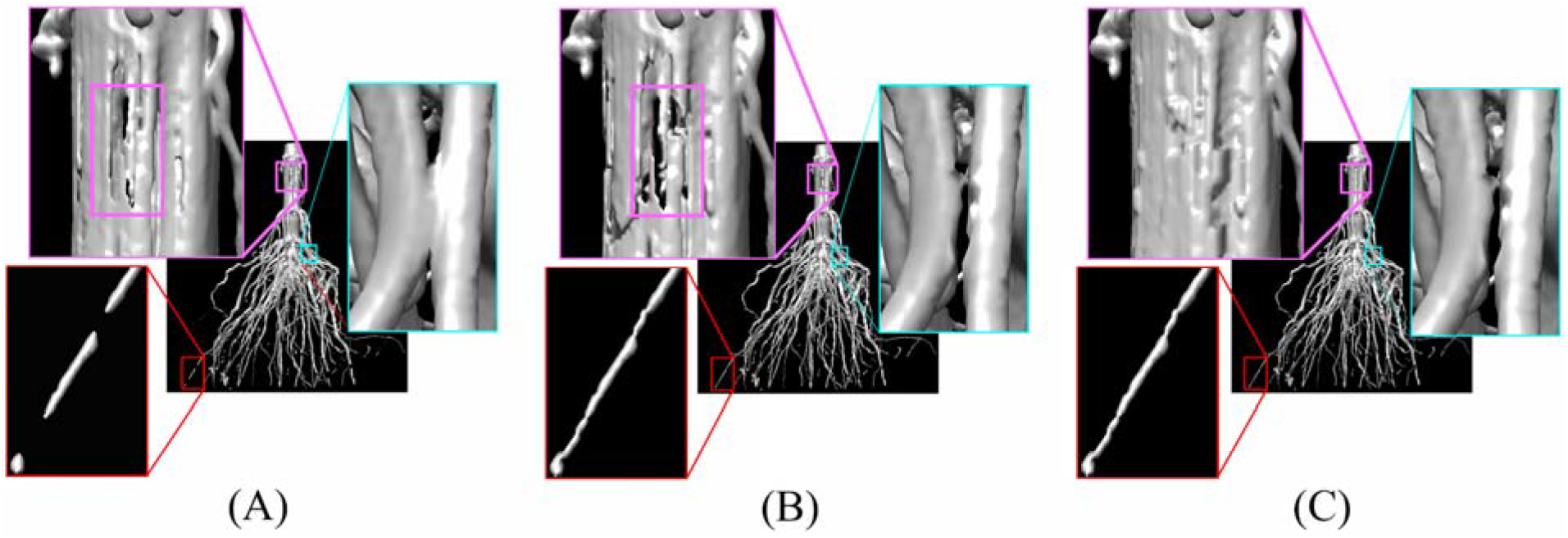
Topological simplification applied to a root image. (A): The input iso-surface at the shape threshold contains numerous topological errors such as disconnections (red box) and loops (cyan box), and tunnels exist on the stem wall (highlighted in the purple box) that connect the hollow interior space to the outside. (B): Applying the algorithm of [38] resolves the disconnections and loops (red and cyan boxes), but the hollow space inside the stem remains connected to the outside through tunnels (highlighted in the purple box). (C): By applying the filling heuristic prior to calling [38], the stem’s interior is filled in without introducing new topological errors.

A common issue in XRT scans of dried roots is that the interior space of the stem and some thick root branches often appears to be hollow. Since our subsequent analysis of the hierarchy relies on a skeleton representation of the root architecture, we need to fill in such hollow space so that the resulting skeleton captures the solid cylindrical shape of the roots (instead of only the walls). However, identifying such interior space is not straight-forward. In the ideal scenario, at the shape threshold, the wall of the stem or root branch completely separates the interior space from the outside space, making the interior space a topological void and hence easy to detect. But oftentimes the iso-surface at the shape threshold exhibits “tunnels” on the wall of the stem or branch (see Figure 5A, top-left purple box), which connect the interior space with the outside and making it impossible to identify as a topological feature. Applying the algorithm of [38] does not fill in the hollow space. On the contrary, it may merge nearby small tunnels into larger tunnels to reduce the number of topological loops (see Figure 5B, top-left).

We develop a simple heuristic to identify and fill the hollow interior spaces. Our assumption, based on observation of our data, is that such spaces usually become topological voids at the lower, *neighborhood* threshold (see Figure 6A). These voids, however, are usually smaller than the hollow space at the shape threshold *V_n_*, and hence need to be expanded before filling in. To do so, we first erode the set of all voxels above the neighborhood threshold, noted as, onto the set of voxels above the shape threshold, noted as *V_s_*, while preserving the topology of *V_n_*. The erosion is prioritized by the intensity, such that voxels having lower intensity are eroded earlier. The eroded voxel set, denoted as *V_n_*’, consists of *V_s_* and a minimal set of voxels needed to achieve the topology of *V_n_*, denoted as *V_f_* (colored red in Figure 6B). We then fill each void of *V_n_*’, as well as those voxels in *V_f_* adjacent to these voids (see Figure 6C). Since filling may introduce additional topological errors, such as loops or new voids, we call the heuristic prior to applying the algorithm of [38]. This produces a topologically simple root shape with filled stems and branches (see Figure 6C).

**Fig. 6.**
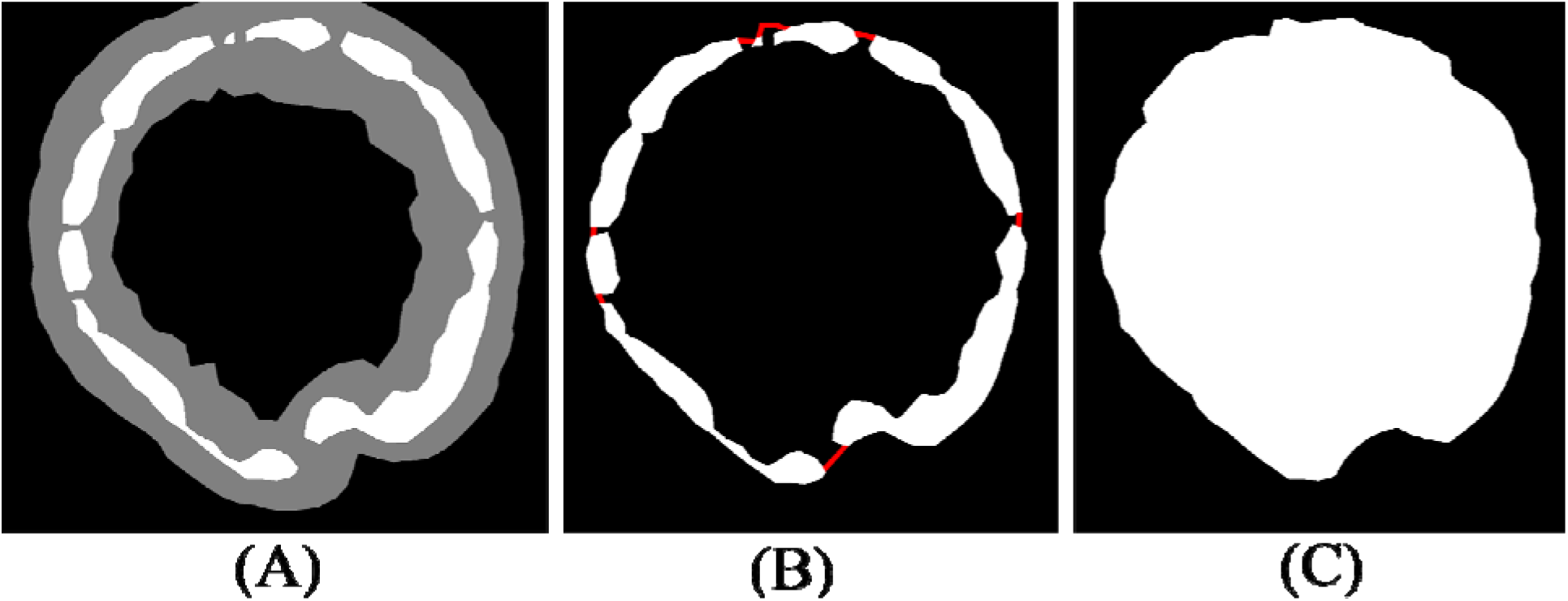
Filling the hollow space inside the stem and thick branches. (A) A cross section of a stem, showing voxels at or above the shape threshold (white, denoted by *V_s_*) and voxels at or above the neighborhood threshold (gray, denoted by *V_n_*). Note that the hollow interior of the stem is connected to the outside space in *V_s_*, but it forms a void disconnected from the outside in *V_n_*. (B) The result of a topology-preserving erosion of *V_n_* onto *V_s_*, denoted by 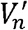, which consists of *V_s_* (white) and additional voxels (red) needed to retain the topology of *V_n_*. (C) The voids of 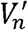, as well as any voxels that are in 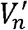 but not in *V_s_* (denoted *V_f_*) are filled in.

Not all topological errors will be removed at the end of this step. In particular, the algorithm of will retain those topological features (e.g., loops) that exist on all three thresholds (shape, kernel, and neighborhood). Removing these larger-scale errors often requires more careful analysis of the root architecture, for example, to prevent breaking a root in the middle. We shall address these remaining issues in the next step of our pipeline and with the help of a geometric skeleton.

### Skeletonization

To obtain the root hierarchy and associated traits, our method relies on a representation of the root system as a curve skeleton - a connected set of curves lying in the center of the root branches and capturing the branching structure. While there are many algorithms for computing skeletons of 3D shapes, we adopt the methods of [36, 37] because of their scalability to high-resolution iso-surfaces, robustness to noise, and the final representation as an embedded graph (with vertices and edges) which is more convenient for connectivity analysis than voxel-based skeletons. These methods have been previously adopted in skeleton-based phenotyping of sorghum panicles [9]. Specifically, given the topologically simplified iso-surface produced by the previous step, [37] computes a 2-dimensional centered structure known as the medial axis, and [36] further reduces the medial axis to 1c-dimensional curves. The resulting skeleton is also associated with shape attributes, including the thickness and the width of the tubular cross-sections, which are useful in our subsequent analysis. An example is shown in Figure 4C (also in Figure 7A).

**Fig. 7.**
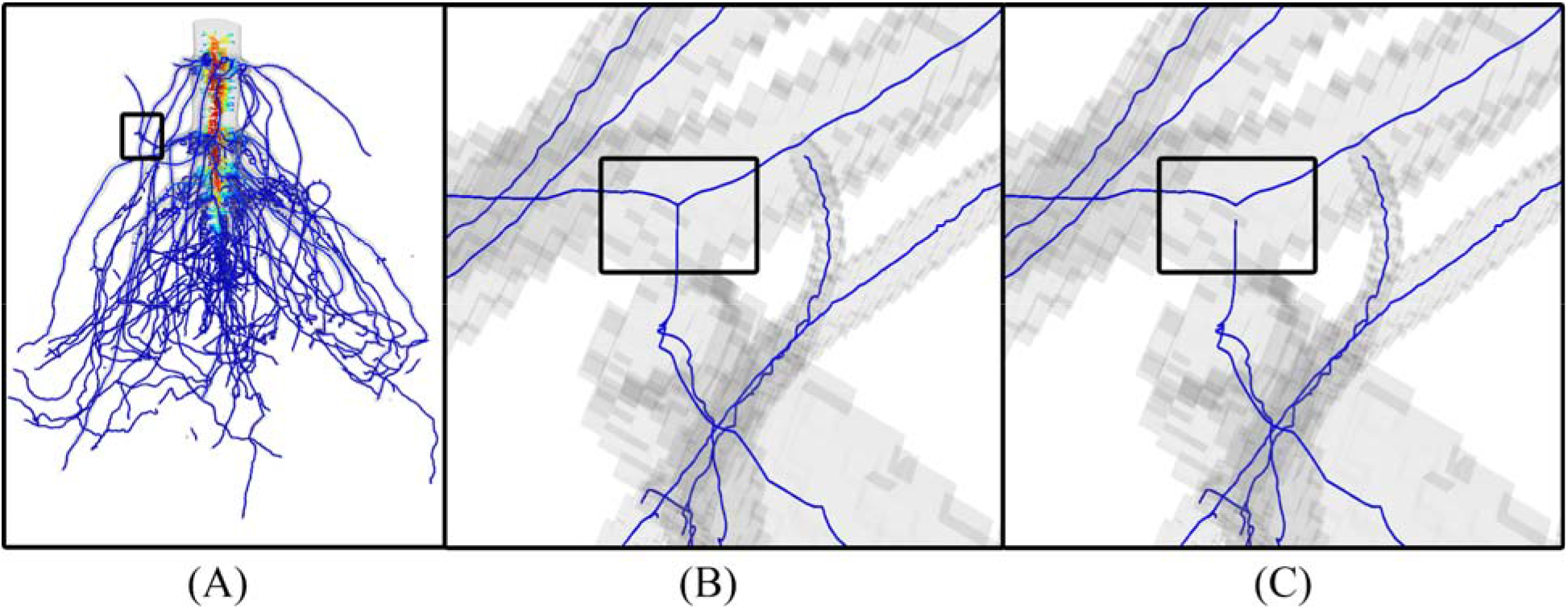
Removing cycles on the curve skeleton. (A) A curve skeleton computed by the methods of [36, 37], where color indicates the thickness of the roots (redder curves lie in thicker roots). (B) A skeleton junction (in the boxed region in (A)) caused by merging between two roots in the iso-surface, which leads to a cycle in the skeleton. (C) The cycle is removed by detaching a skeleton segment from the junction.

As mentioned earlier, some topological errors remain after topological simplification. These errors manifest as disconnected components and cycles (a path of edges which begin and end at the same vertex) on the curve skeleton. To reduce the number of components to one, we simply take the largest connected component on the skeleton. To remove cycles in that component, we observe that cycles are usually caused by merging of distinct roots, which take place at junctions (vertices with degree three or higher) on the skeletons (Figure 7B). Our goal is to identify junctions that correspond to merging between roots, as opposed to natural branching of roots, and resolve the cycles by detaching skeleton segments from such junctions (Figure 7C). This is achieved by computing the minimum spanning tree (MST) on a weighted graph, as described below.

As we are primarily concerned with the junctions of the skeleton, we construct an abstract graph where each node represents either a junction vertex or a continuous skeleton segment between junctions (Figure 8A, 8B). A graph edge connects a node representing a skeleton junction with another node representing a skeleton segment incident to the junction. Removing a graph edge corresponds to detaching a skeleton segment from a junction (Figure 8C, 8D).

**Fig. 8.**
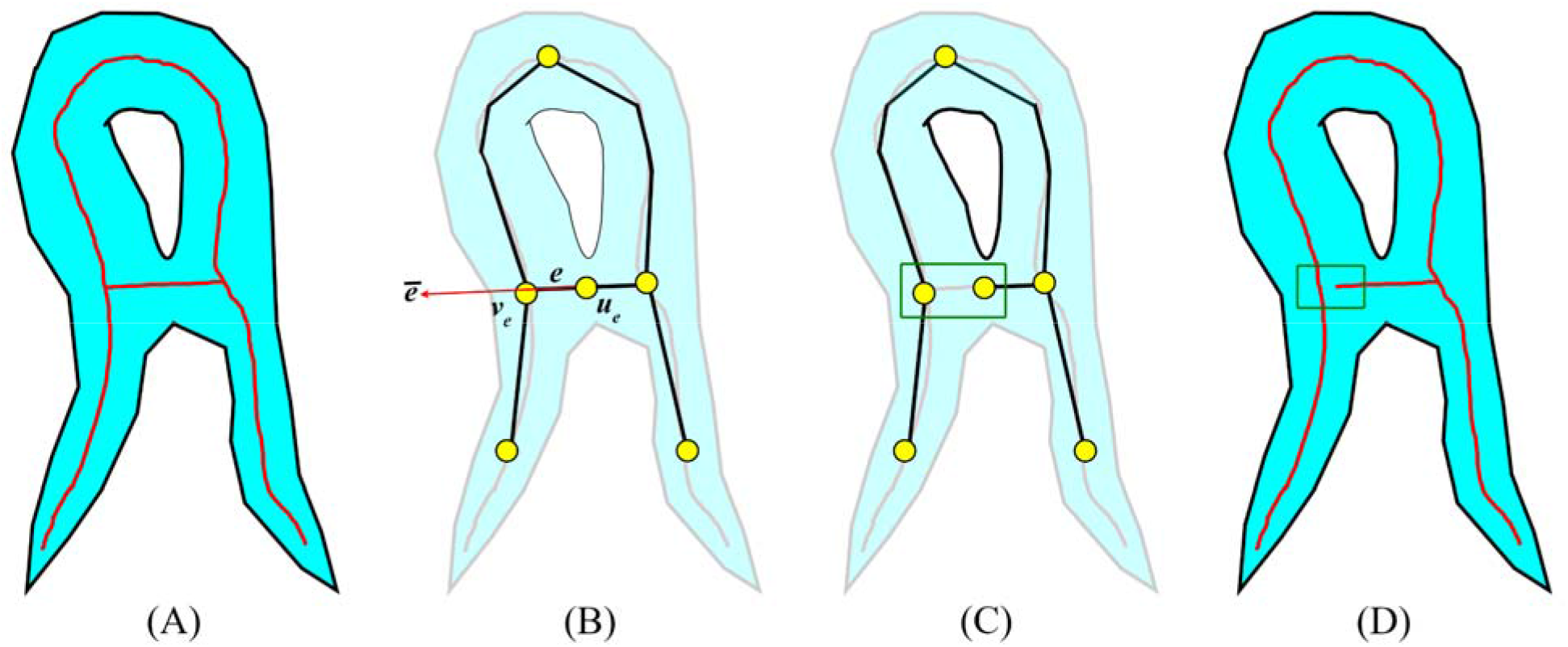
Illustration of our cycle-breaking algorithm. (A) A curve skeleton (red) of a synthetic root system (blue) that contains a cycle. (B) A graph that represents each skeleton junction or segment by a node (yellow circle) and connects two nodes representing a junction and an incident segment by an edge (black line). *e* is a graph edge, *v_e_* is the node that represents a skeleton junction, *u_e_* the other node of *e* which represents a skeleton segment, and *<u>e</u>*is the unit vector of the tangent direction of the skeleton segment represented by node*u_e_* towards the junction node *v_e_*. (C) The MST of the graph removes the cycle by excluding an edge (green box). (D) The corresponding skeleton segment is detached from the skeleton junction.

The MST is the subset of edges such that all graph nodes remain connected and the sum of edge weights is minimal. An MST is always free of cycles. We define the edge weights such that a lower weight implies a higher likelihood that a skeleton segment should be attached to a junction. Our weight definition is motivated by the observation that a merging between two distinct roots often results in a “T” junction on the skeleton where, among all skeleton segments incident to the junction, one segment (which should be detached) is close to being orthogonal to the other segments (see Figure 8B). In contrast, a typical branching of the root usually results in a “Y” junction on the skeleton where each skeleton segment at the junction forms an obtuse angle with at least one other segment. However, branching close to the stem of the root could be more like “T” than “Y”, due to the large difference in thickness between the main and offspring roots. Our weight encourages “Y” junctions while preventing detachment close to the stem.

Specifically, let *e* be a graph edge, *v*_e_ the node of the edge *e* that represents a skeleton junction, *u*_e_ the other node of *e* which represents a skeleton segment, and *E(v)* the set of edges incident to node *v*. We denote by *ē* the unit vector of the tangent direction of the skeleton segment represented by node *u*_e_ *towards* the junction node *v*_e_. We first define the angle deviation term *w*_*angle*_ as:

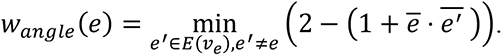

This term reaches the minimal value of 0 when there is some edge *e′* incident to *v*_e_ that has the same direction as *e*, and it attains the maximal value of 2 when all other edges incident to *v*_e_ have the opposite direction as *e*. Next, we define *w*_disc_(*e*) as the shortest distance along the skeleton between the skeleton junction represented by ve and any skeleton vertex in the stem. Here, we extract the stem from the skeleton using the heuristic reported in [9]. The heuristic starts from a subset of the skeleton whose thickness measure is above a threshold (since the stem is usually the thickest portion of the maize root system) and extracts the longest simple path from the subset. Finally, the edge weight is defined as:

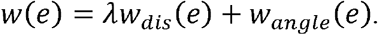

Here *λ* is a balancing parameter, which is set to 0.05 in our experiments.

### Inferring hierarchy

A key feature of TopoRoot is computing the hierarchy of the maize root system consisting of the stem, nodal roots, and lateral roots at different levels. Given the cycle-free skeleton computed from the previous steps, we next label each vertex of the skeleton as either part of the stem path, the stem region, a nodal root, or a lateral root at a specific level. As described in the previous step, the stem path is identified using the thickness-based heuristic of [9]. The stem region is then defined as the set of skeleton edges within a cylindrical region around the skeleton path, where the radius of the cylinder varies along the stem path and is set to be 1.6 times the thickness measure stored on the path vertices. In Figure 4D, the stem path and stem region are colored dark blue.

To label the remaining skeleton edges by the root hierarchy (e.g., nodal roots, 1st-order lateral roots, 2nd-order lateral roots, etc.), we make the following two assumptions. First, as assumed in previous works including DynamicRoots, roots at lower levels of the hierarchy are generally longer. For example, nodal roots are generally longer than lateral roots, and 1st-order lateral roots are generally longer than 2nd-order lateral roots, and so on. Second, the maximum number of hierarchy levels in a root system is generally kept low. With these assumptions, we developed a labelling heuristic that minimizes the maximal level of hierarchy in the root system while encouraging longer roots to be at lower levels.

Our heuristic proceeds in two stages, a bottom-up traversal of the skeleton and then a top-down traversal. They are illustrated in Figure 9 using a cartoon example. We start with a skeleton labelled only by the stem region (Figure 9A). Since the skeleton has no cycles, it is a “tree”. We consider the stem region as the “root” of this tree, and this induces a partial ordering of the skeleton segments such that each skeleton junction (outside the stem region) is incident to exactly one *parent* segment and zero or more *children* segments. The first stage of the heuristic computes, at each skeleton junction, the association between the parent skeleton segment with one of the children segments as the continuation of the same root (see arrows in Figure 9B). The association is computed by visiting the skeleton segments from the leaves of the skeleton towards the stem and updating a depth *d(b)* (numbers in Figure 9B) and a distance *l(b)* at each visited segment *b*, as follows. First, for each segment *b* incident to a leaf vertex of the skeleton, we set *d(b)*=0 and *l(b)* as the length of the segment *b*. We then iteratively visit parent segments whose children segments have already been visited. For a parent segment *b* whose children are *b_1_*, …., *b_n_*, we associate *b* with the child *b*_*i*_ that has the maximal depth *d(b_i_)*. If multiple children have the same maximal depth, *b* is associated with the *b*_*i*_ with maximal length I(b_*i*_). We then set *d(b)=d(b_*i*_)+1* and *l(b)* to be *l(b_*i*_)* plus the length of segment *b*. In the second stage, we visit all segments from the stem to the leaves and assign the hierarchy levels. We assign each segment attached to the stem region a hierarchy level of 1 (i.e., nodal roots). For each parent segment assigned with level k, we assign level *k* to the child segment associated with the parent (computed from the first stage) and level *k* +1 to all other children segments. The resulting hierarchy labelling is shown in Figure 9C both as numbers and the heat color (warmer colors have higher levels).

**Fig. 9.**
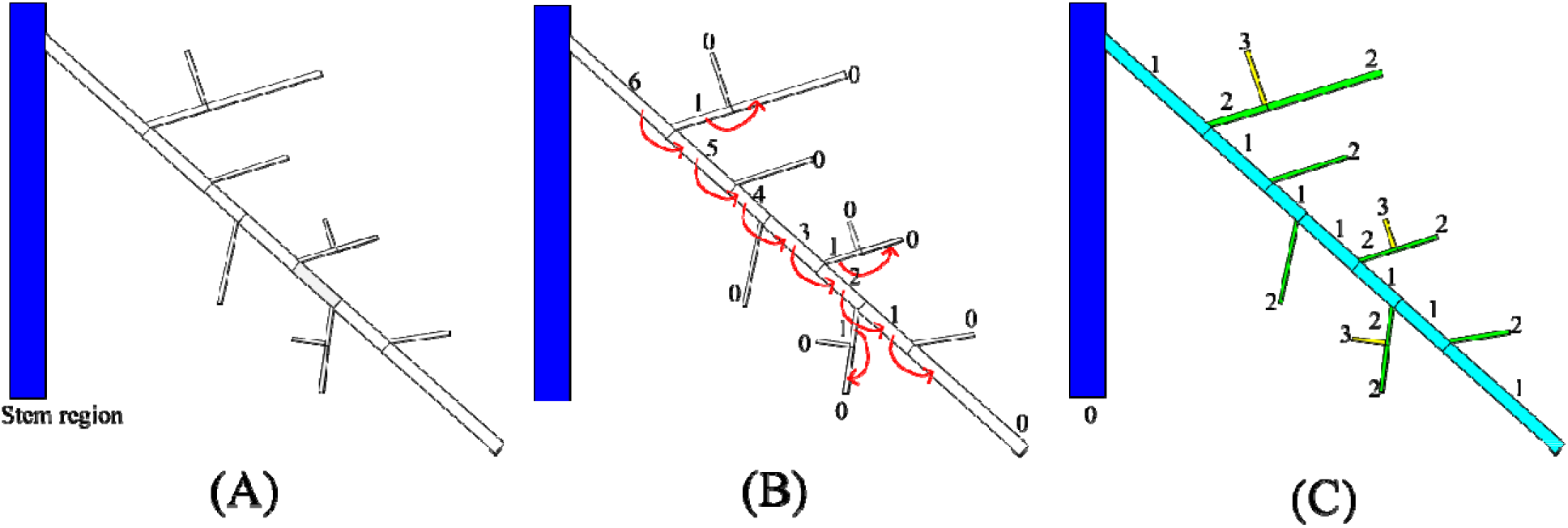
Heuristic for inferring hierarchy on a cycle-free skeleton. (A) The input skeleton with only the stem region labelled (blue). (B) The first stage associates each parent segment with one of its children segments (indicated by arrows). The numbers are the depth stored at each segment, an intermediate quantity used to determine the parent-child association. (C) The second stage assigns hierarchy levels to each skeleton segment based on the parent-child association.

### Computing traits

Given the skeleton and the hierarchy labelling, TopoRoot computes a suite of coarse-grained and fine-grained traits of the root system. Like existing works (e.g. [33]), we compute global traits which are aggregated over all the hierarchy levels, including the total root length, number of roots, and average root length. For fine-grained traits, for each hierarchy level (e.g., nodal roots, 1st-order lateral roots, 2nd-order lateral roots, etc.), we compute the root count, average and total root length, average root tortuosity, average root thickness, average number of children, and the average emergence, midpoint, and tip angle. We also report the length and thickness of the stem. Some of these traits, such as stem length and per-level angle traits, have not been previously reported by existing tools (including DynamicRoots [32]). Details on how each of these traits is computed can be found in supplementary table 1.

**Table 1:** Reporting the accuracy of TopoRoot for stem traits.

| Trait | $\leq e$ | TopoRoot (%) |
| --- | --- | --- |
| Stem length | 0.04 | 6.9 ($\sigma=6.9$) |
| | 0.08 | 7.7 ($\sigma=7.3$) |
| Stem thickness | 0.04 | 11.9 ($\sigma=4.3$) |
| | 0.08 | 15.3 ($\sigma=5.8$) |

Each entry gives the average error (and standard deviation $\sigma$ ) for our method across all simulated models and across all noise levels up to $e=0.04$ cm and 0.08 cm.

## Results

We validated TopoRoot on the aforementioned dataset and compared it with some previous tools, including DynamicRoots [32] (for both global and fine-grained traits) and a 3D version of GiaRoots [4] first published in [33] (for global traits only). We used default parameters for both DynamicRoots and GiaRoots. DynamicRoots requires a “seed” voxel, which we set as a random voxel from the top slice. We ran DynamicRoots and GiaRoots on the thresholded images at the shape threshold. To illustrate the effect of topological errors on these tools, we extend each tool by first performing the topological simplification step of TopoRoot. We call the extended tools DynamicRoots+ and GiaRoots+ respectively. TopoRoot is implemented in C++, and all experiments were run on a Windows 10 machine with an Intel(R) Core(TM) i9-10900X Processor @ 3.70 GHz and 64.0 GB of memory (RAM).

### Excavated root crowns

First, we visually compare the root hierarchies computed by TopoRoot, DynamicRoots, and DynamicRoots+ for a randomly picked root crown (Figure 10). DynamicRoots produces a point cloud where each point represents an input voxel and is labelled by its hierarchy level (0, 1, 2, etc.). As explained in [32], the hierarchy levels produced by DynamicRoots reflect the geometric branching structure and may not map well to the biological hierarchy (e.g., stem, nodal roots, lateral roots, etc.). For this reason, as well as the inaccuracy in determining hierarchy without a dense time series and the different morphology of mature root crowns in our dataset from those of seedlings grown in a gel environment (for which DynamicRoots was designed), DynamicRoots tends to produce significant mis-classifications of root branches in our dataset.

**Fig. 10.**
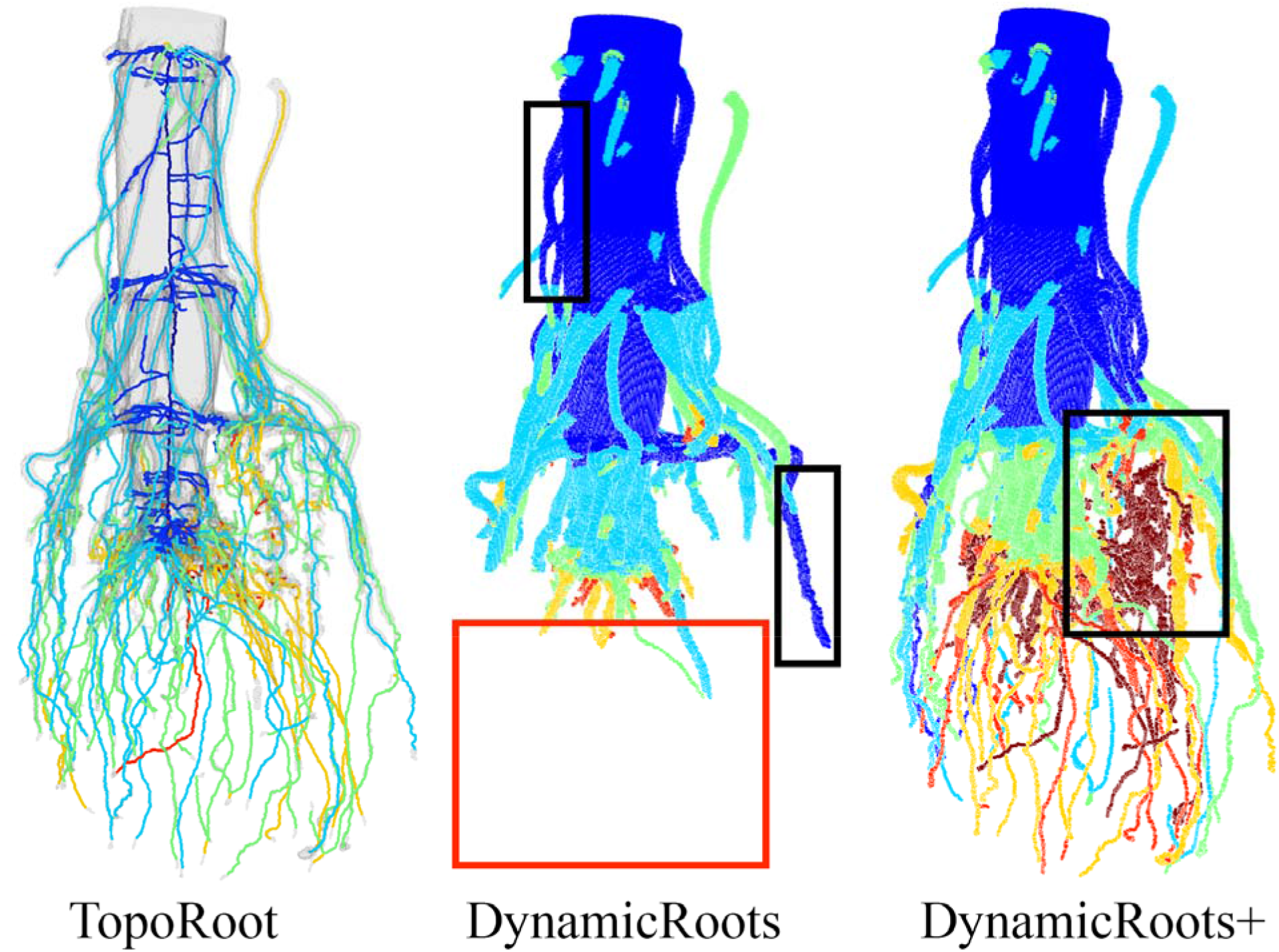
Visual comparison of root hierarchies computed by TopoRoot, DynamicRoots and DynamicRoots+ from the X-ray CT scan of an excavated maize root crown. Hierarchy levels are colored as follows: 0 (stem): dark blue, 1 (nodal roots): light blue, 2 (1st-order lateral roots): green, 3 (2nd-order lateral roots): orange, 4 (3rd-order lateral roots): red, ≥ 5: dark red. Black boxes highlight incorrect levels obtained by DynamicRoots and DynamicRoots+, and the red box highlights a missing component in DynamicRoots.

For example, the level-0 roots obtained by DynamicRoots include not only the stem but quite a few nodal roots (black boxes). These errors propagate to higher-level roots, which fork from lower-level roots. In addition, since the thresholded root image at the shape threshold contains many disconnected components, and because DynamicRoots only considered one connected component, a significant portion of the root is missing in the analysis of DynamicRoots (red box). Although performing the topological simplification step in TopoRoot allows DynamicRoots+ to recover a more complete root system, mis-labelling of hierarchy levels remains (black boxes). In contrast, TopoRoot produces a more visually plausible hierarchy separating the stem (region), nodal roots, and lateral roots.

Next, we quantitatively validate the nodal root counts computed by various tools against the hand measurements. Figure 11 plots the correlation between the hand-measured nodal root counts and those computed by TopoRoot (A), DynamicRoots (B) and DynamicRoots+ (C) for all 45 samples. Based on visual inspection (Figure 10), we consider the level 1 branches as nodal roots for DynamicRoots+. For each sample we computed the relative error by taking the absolute difference of TopoRoot’s nodal root count and the hand measurement, and finding the percentage of this difference relative to the hand measurements. We compute the mean and standard deviation (*σ*) of this relative error for TopoRoot, DynamicRoots, and DynamicRoots+. TopoRoot exhibits a much lower relative error (mean=8.3%, *σ* = 5.6%) and higher correlation (Pearson’s coefficient=0.951) than either DynamicRoots (159.5% mean error with *σ* = 190.0%, Pearson’s coefficient=−0.0452) or DynamicRoots+ (235.4% mean error with *σ* = 244.4%, Pearson’s coefficient=−0.160). The significant over-counting of DynamicRoots+ is mostly caused by the mislabeling of nodal roots as level-0 roots, as explained above, which leads to many lateral roots being labelled as level-1 roots. The over-counting also increases with the size of the root system. Furthermore, both the nodal root counts computed by TopoRoot and the hand measurements exhibited a significant difference between the mutant and wild type samples, as measured by the independent two-sided Wilcoxon rank sum test (p=0.000130 for TopoRoot, p=0.00349 for hand measurements). Neither DynamicRoots (p=0.126) nor DynamicRoots+ (p=0.0199) showed a significant difference between the mutant and wild-type, and both had negative Pearson coefficients. This shows that TopoRoot can perform better for differentiating the root system architecture between these two varieties than can DynamicRoots.

**Fig. 11.**
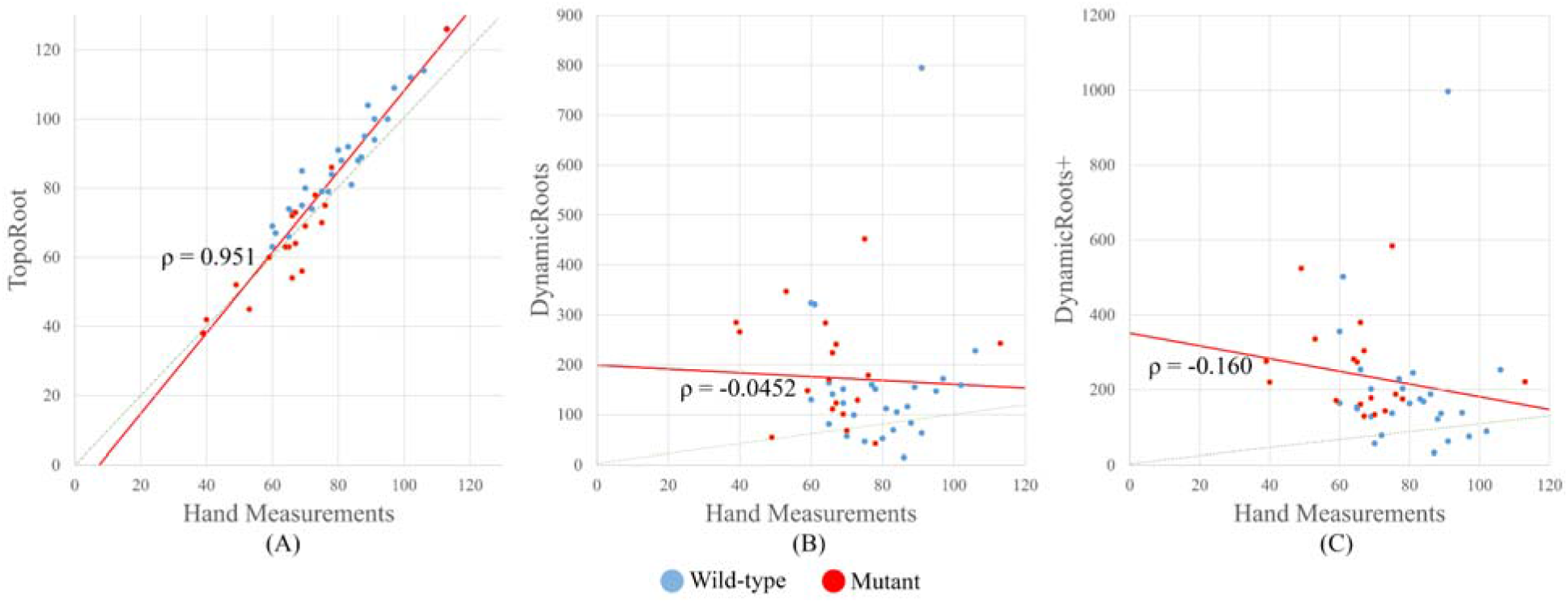
Correlation of nodal root count between hand and computational measurements. (A) Correlation between TopoRoot and hand measurement. (B) Correlation between DynamicRoots and hand measurement. (C) Correlation between DynamicRoots+ and hand measurement. Each dot in the graph represents one of the 45 samples. Blue and red dots indicate wild type and mutant samples, respectively. The regression line is in red, while the dashed green line indicates the ideal correspondence between the hand and computational measurements. The correlation coefficient ρ is indicated in each plot.

Since we do not have hand measurements of other fine-grained traits for this data set, we perform an indirect evaluation by assessing the ability of each trait computed by TopoRoot in differentiating the mutant and wild type samples. As shown in Figure 12, 12 out of the 23 fine-grained traits computed by TopoRoot (reported up to the hierarchy level of the first-order lateral roots) exhibit a significant difference between the two genotypes (p<0.01). Compared to the wild type, the mutants generally have fewer and shorter thinner roots at each hierarchy, whereas the various angle measures are greater. These fine-grained trait differences offer a more comprehensive analysis of phenotypic differences caused by the mutation which better characterizes gene function and may lead to novel biological investigations.

**Fig. 12.**
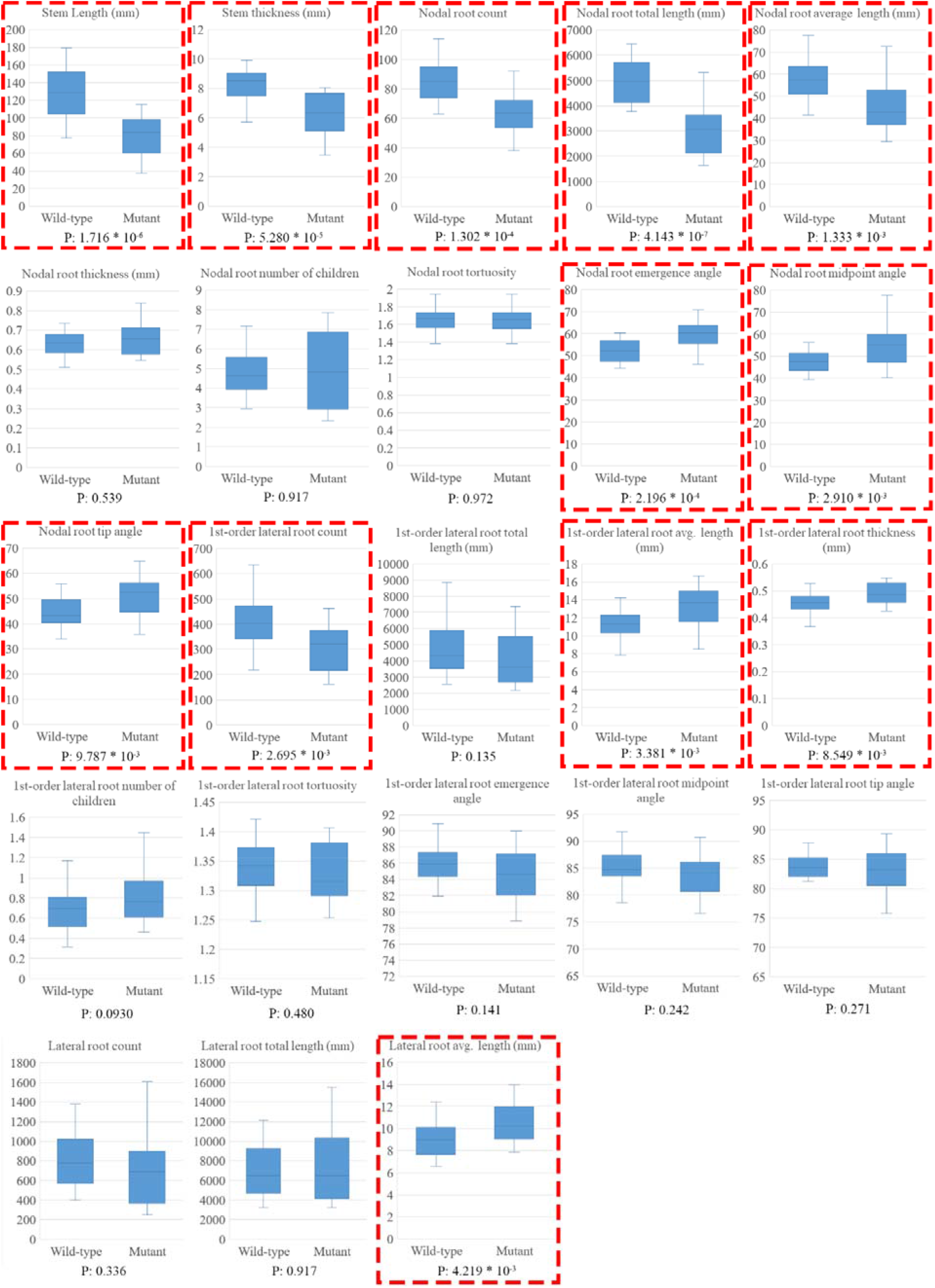
Fine-grained traits computed by TopoRoot for wild type and mutant samples. Each boxplot shows distribution bars for one trait over all 27 wild type samples and 18 mutants. Traits with p<0.01, as measured by the independent two-sided Wilcoxon rank sum test, are highlighted in red boxes.

### Simulated Roots

We start with a visual comparison of the results of TopoRoot, DynamicRoots+ and GiaRoots+ on one of the simulated root systems. This root system is simulated to be 34 days old, with five whorls, 34 nodal roots, and a lateral root branching frequency between 0.3 - 0.7 cm / branch. Figure 13 visualizes the root hierarchies produced by TopoRoots and DynamicRoots+ as well as the voxelized skeletons produced by GiaRoots+ at three different noise levels (0, 0.04cm, 0.08cm). As the noise level increases, the thresholded image at the shape threshold becomes more disconnected and contaminated by topological errors (see Figure 13). Accordingly, DynamicRoots and GiaRoots miss more root parts, whereas TopoRoot as well as the extended tools, DynamicRoots+ and GiaRoots+, retain much of the root shape. Observe that, similar to the real roots dataset, the hierarchies produced by DynamicRoots+ incorrectly label many nodal roots as level 0 (black boxes). In contrast, the hierarchies produced by TopoRoot are more visually plausible.

**Fig. 13.**
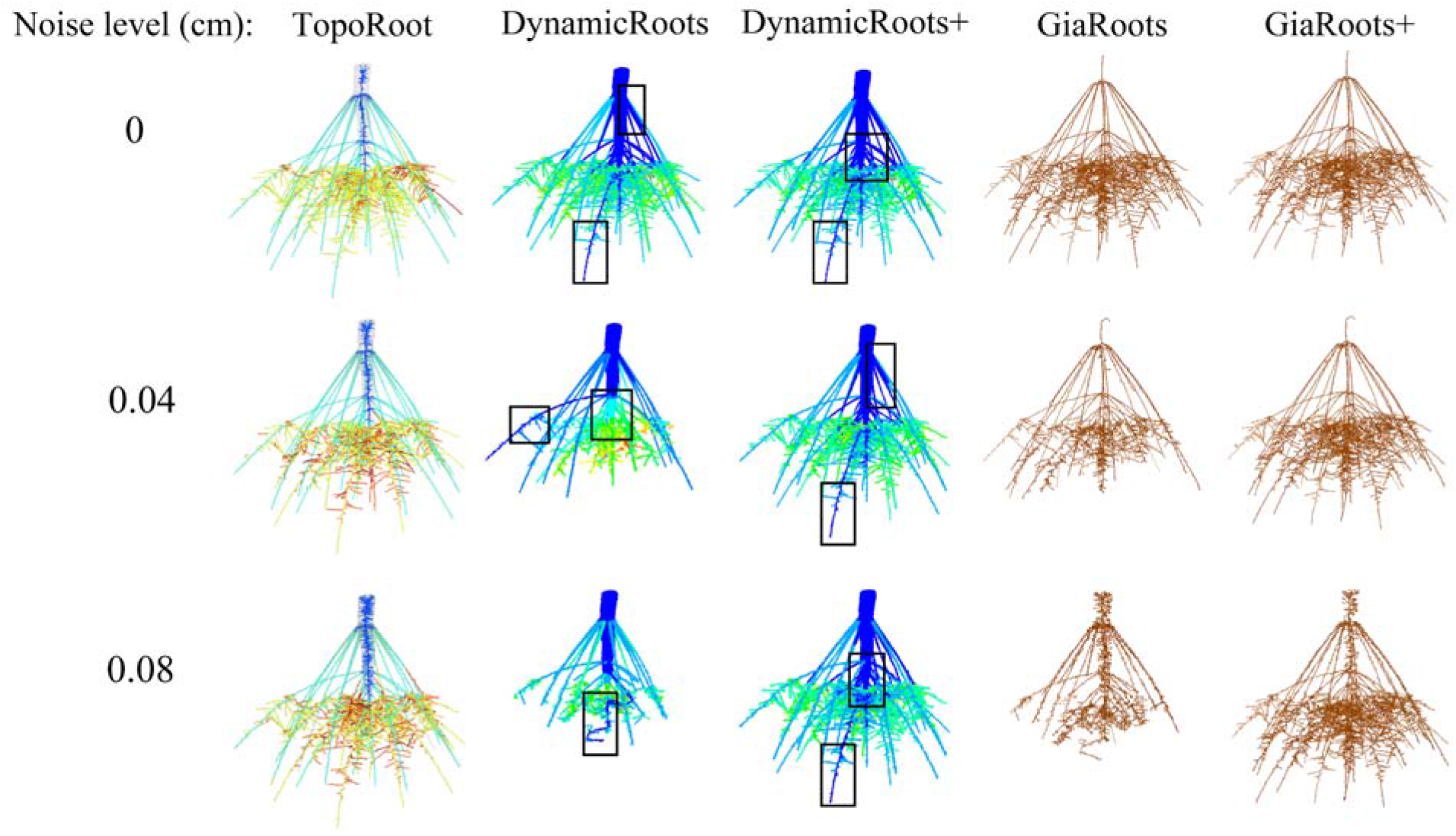
Comparing hierarchies and skeletons computed by different tools from images of a simulated maize root. Three different noise levels are shown. Hierarchy levels 0, 1, 2, 3 and 4 produced by TopoRoot and DynamicRoots+ are colored dark blue, light blue, green, orange, and red. The voxelized skeletons produced by GiaRoots+ are colored brown. Black boxes highlight mislabeling of nodal roots as level 0 roots by DynamicRoots and DynamicRoots+.

We performed a thorough quantitative validation of all traits computed by TopoRoot against the ground truth provided by OpenSimRoot. The relative error for each trait over the entire simulated dataset, as well as the errors of DynamicRoots+ and GiaRoots+ (for global traits only), are reported in Tables 1–4 for noise levels ≤ 0.04 cm and 0.08 cm. In supplementary Figures S1–S4, we take a closer look at the accuracy of TopoRoot and the other tools as a function of the noise level of the input images. These figures plot the relative errors of the five tools in computing the stem traits (Figure S1), the nodal root traits (Figure S2), the lateral root traits (Figure S3), and the global traits (Figure S4) as the noise level increases. Roots less than one voxel long in the ground truth model were ignored in our analysis. We compare and analyze the accuracy of TopoRoot across each category of traits below. In general, we observe that higher image noise leads to larger mean errors and/or greater variance by TopoRoot. For most of the traits, TopoRoot maintains a lower error than other tools across all noise levels.

As the base of the hierarchy, the stem traits are among the most accurate. As the noise increases, portions of the stem region are lost, resulting in a thinner stem (Figure S1). Increased noise also causes the stem to wiggle more in the direction perpendicular to its main path, resulting in an increased stem length.

The nodal root traits (Table 2) are the most accurate for the root count (8.3% up to *e*=0.04, 10.3% up to *e*=0.08) and emergence and midpoint angles (5.7/7.1% up to *e*=0.04, 9.1/9.5% up to *e*=0.08). The lowest accuracy is seen for the number of children (40.0% up to *e*=0.04, 48.6% up to *e*=0.08) and thickness (39.2% up to *e*=0.04, and 38.2% up to *e*=0.08). These errors are due to misclassifications when the nodal root becomes entangled with higher-order lateral roots. TopoRoot slightly underestimates the average and total length due to faulty cycle breaking and misclassification errors in portions of nodal roots further away from the stem. TopoRoot’s error in nodal root tortuosity is higher than that of DynamicRoots for two reasons. First, the ground truth tortuosity is close to 1 (only for the simulated data, but not for the real maize roots), and DynamicRoots coincidentally produces values close to this because it mistakes many shorter lateral roots as nodal roots, as evidenced by its much shorter average nodal root length and the black boxes of Figure 13. Second, nodal roots sometimes are misclassified by TopoRoot closer to their tips due to the large number of intersections between roots of different hierarchy levels, resulting in excessive winding. TopoRoot slightly overestimates angle measurements due to misclassification errors further away from the stem which bend the detected paths sideways; these explain the errors in the tip angle measurements.

**Table 2:** Comparing TopoRoot to DynamicRoots and DynamicRoots+ for nodal root traits.

| Trait | $\leq e$ | TopoRoot (%) | DynamicRoots (%) | DynamicRoots+ (%) |
| --- | --- | --- | --- | --- |
| Nodal root count | 0.04 | **8.3** ($\sigma=8.2$) | 392.9 ($\sigma=423.5$) | 486.2 ($\sigma=426.6$) |
| | 0.08 | **10.3** ($\sigma=10.9$) | 263.9 ($\sigma=367.0$) | 455.2 ($\sigma=400.0$) |
| Nodal root total length | 0.04 | **22.4** ($\sigma=13.1$) | 46.4 ($\sigma=17.7$) | 38.3 ($\sigma=14.3$) |
| | 0.08 | **23.9** ($\sigma=13.7$) | 56.9 ($\sigma=19.7$) | 40.7 ($\sigma=14.6$) |
| Nodal root average length | 0.04 | **17.8** ($\sigma=11.2$) | 76.6 ($\sigma=18.8$) | 80.9 ($\sigma=16.1$) |
| | 0.08 | **19.9** ($\sigma=13.1$) | 74.0 ($\sigma=17.4$) | 81.0 ($\sigma=15.4$) |
| Nodal root thickness | 0.04 | 39.2 ($\sigma=27.6$) | **36.6** ($\sigma=33.8$) | 46.2 ($\sigma=115.0$) |
| | 0.08 | **38.2** ($\sigma=26.8$) | 50.1 ($\sigma=150.3$) | 48.2 ($\sigma=109.6$) |
| Nodal root number of children | 0.04 | **36.4** ($\sigma=21.6$) | 87.7 ($\sigma=13.7$) | 89.4.5 ($\sigma=12.5$) |
| | 0.08 | **45.6** ($\sigma=41.0$) | 87.0 ($\sigma=13.1$) | 89.2 ($\sigma=12.4$) |
| Nodal root tortuosity | 0.04 | 32.6 ($\sigma=13.0$) | 2.3 ($\sigma=2.5$) | **2.0** ($\sigma=1.8$) |
| | 0.08 | 37.3 ($\sigma=13.8$) | 4.5 ($\sigma=4.4$) | **2.9** ($\sigma=2.3$) |
| Nodal root emergence angle | 0.04 | **5.7** ($\sigma=11.9$) | N/A | N/A |
| | 0.08 | **7.3** ($\sigma=11.9$) | N/A | N/A |
| Nodal root midpoint angle | 0.04 | **7.1** ($\sigma=14.0$) | N/A | N/A |
| | 0.08 | **9.1** ($\sigma=13.8$) | N/A | N/A |
| Nodal root tip angle | 0.04 | **18.3** ($\sigma=17.7$) | N/A | N/A |
| | 0.08 | **22.7** ($\sigma=22.8$) | N/A | N/A |

Each entry gives the average error (and standard deviation $s$ ) for a method across all simulated models and across all noise levels up to $e=0.04$ cm and 0.08 cm. For each trait, the method with the lowest error is bold-faced.

The errors for the lateral root traits (Table 3) are generally larger than nodal root traits, primarily since the imaging noise has a greater impact on the thinner roots more than the thicker ones. There is a greater underestimation of both the total first-order lateral roots and their total length (Figure S3), due to both the misclassification of the hierarchy levels and the loss of many thin roots in the distance field. On the other hand, the misclassified first-order lateral roots are counted as lateral roots of higher orders, and hence less errors lie in the total lateral root count (22.2% up to e=0.04 and 37.0% up to e=0.08) and lengths (24.3% up to e=0.04 and 24.4% up to e=0.08) over all orders. All methods significantly overestimate the first-order lateral root thickness due to limits in the resolution, but TopoRoot produces the lowest error. The lowest errors are seen in the first-order lateral emergence/midpoint/tip angles (3.4%/4.1%/5.4% up to e=0.04 and 3.0%/4.0%/5.9% up to e=0.08) and tortuosity (4.6% up to e=0.04 and 4.5% up to e=0.08).

**Table 3:**
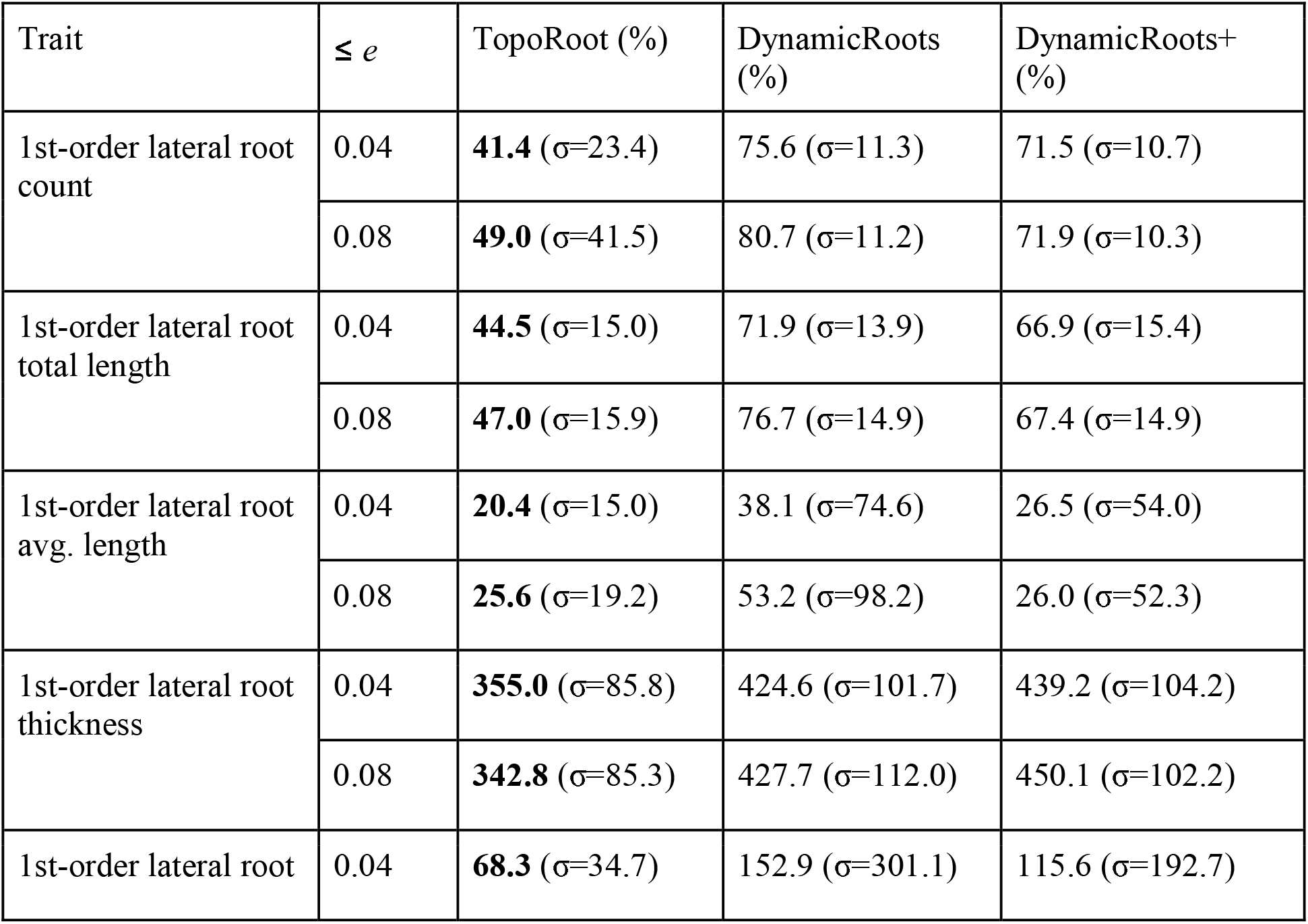
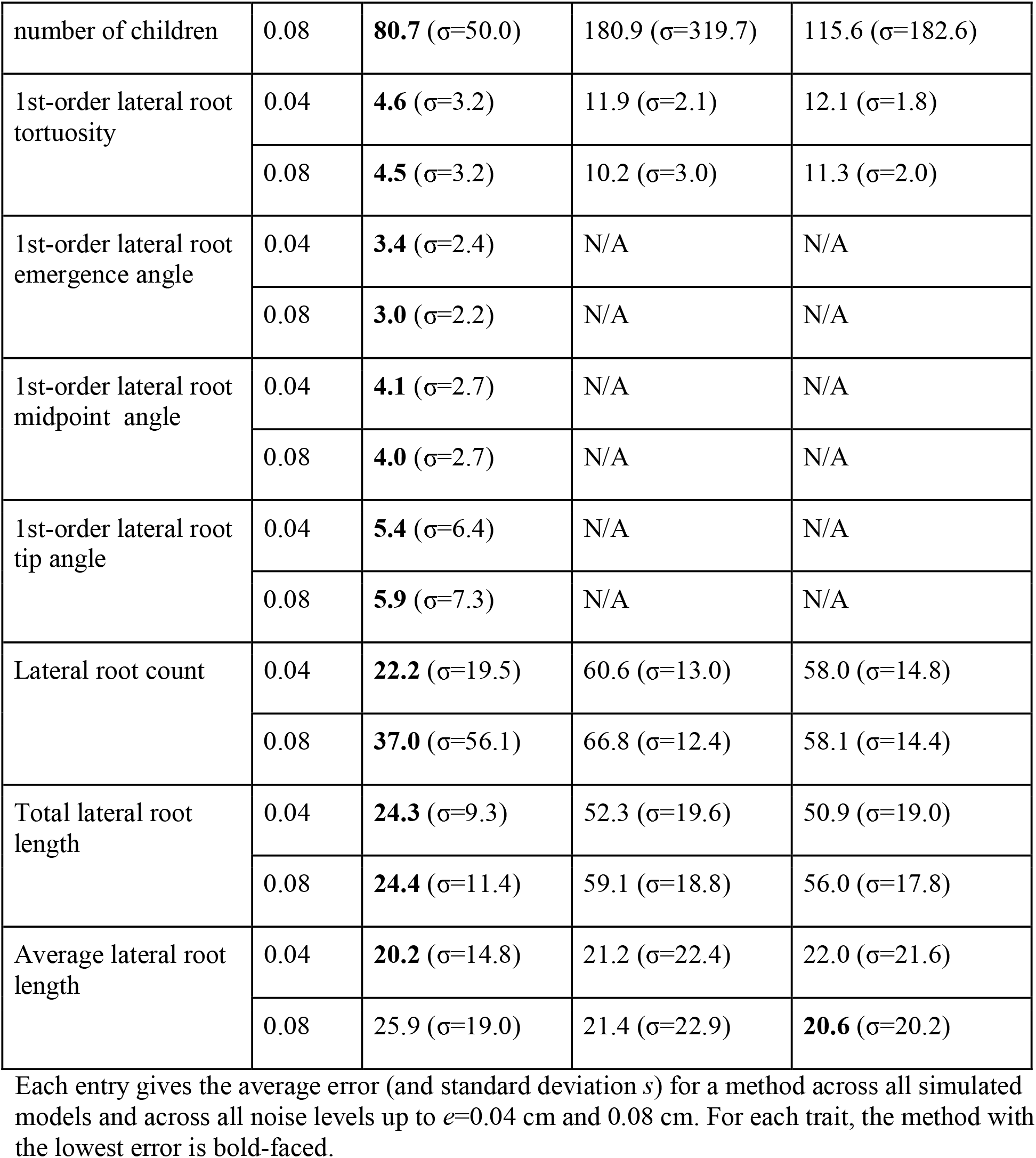
Comparing TopoRoot to DynamicRoots and DynamicRoots+ for lateral root traits.

Finally, combining nodal and lateral roots, TopoRoot produces on average 35.4% error (21.5% up to e=0.04) in the total root count and 25.4% error (25.0% up to e=0.04) in the total root length, which are much lower than DynamicRoots and GiaRoots (Table 4). Note that both DynamicRoots and GiaRoots significantly underestimate the root count and total length, even after topological simplification, and the amount of underestimation generally increases with the level of noise (Figure S4). The only global trait that TopoRoot does not have the lowest error is the average length, due to a combination of DynamicRoots being coincidentally closer due to its underestimation of both the total length and number of roots, and TopoRoot having an excessive number of roots at higher noise levels. These are the same reasons why the two methods have similar lateral root average length errors.

**Table 4:** Comparing TopoRoot to DynamicRoots, DynamicRoots+, GiaRoots, and GiaRoots+ for global traits.

| Trait | $\leq e$ | TopoRoot (%) | DynamicRoots (%) | DynamicRoots+ (%) | GiARoots (%) | GiARoots+ (%) |
| --- | --- | --- | --- | --- | --- | --- |
| Total root count | 0.04 | **21.5** ($\sigma=18.4$) | 53.7 ($\sigma=11.1$) | 49.3 ($\sigma=11.3$) | 57.9 ($\sigma=13.7$) | 56.8 ($\sigma=13.7$) |
| | 0.08 | **35.4** ($\sigma=51.8$) | 60.8 ($\sigma=12.7$) | 49.7 ($\sigma=11.3$) | 45.7 ($\sigma=21.1$) | 57.5 ($\sigma=13.2$) |
| Average root length | 0.04 | 18.4 ($\sigma=13.5$) | 9.1 ($\sigma=11.8$) | **9.7** ($\sigma=9.8$) | 42.6 ($\sigma=33.5$) | 50.2 ($\sigma=35.3$) |
| | 0.08 | 24.7($\sigma=18.8$) | **9.9** ($\sigma=12.5$) | 13.3 ($\sigma=12.2$) | 40.8 ($\sigma=27.4$) | 55.6 ($\sigma=34.2$) |
| Total length | 0.04 | **25.0** ($\sigma=8.5$) | 52.2 ($\sigma=13.9$) | 48.7 ($\sigma=13.1$) | 46.0 ($\sigma=12.1$) | 40.1 ($\sigma=12.1$) |
| | 0.08 | **25.4** ($\sigma=9.7$) | 59.6 ($\sigma=14.5$) | 52.8 ($\sigma=12.4$) | 45.7 ($\sigma=19.6$) | 38.2 ($\sigma=13.1$) |

Each entry gives the average error (and standard deviation $s$ ) for a method across all simulated models and across all noise levels up to $e=0.04$ cm and 0.08 cm. For each trait, the method with the lowest error is bold-faced.

## Discussion

A gap exists in the phenotypic measure of root system architecture between fine-grained analyses that can be conducted on entire seedling root systems in laboratory settings, and much coarser global analyses available to field researchers. Since root systems are an emergent property of their many hundreds, thousands, or tens of thousands of constituent roots, this gap is a major hindrance to a comprehensive understanding of root system development, environmental interaction, and the genetics that influence these processes. In previous work, we showed that when global 3D analysis of field excavated maize root crowns was compared to 3D seedling analysis in gellan gum, genetically encoded differences were consistent despite major differences in developmental stage and the growth environment [17]. Whereas DynamicRoots was previously developed for fine scale measurements of 3D seedling root systems containing dozens to hundreds of roots, no similar tool existed for more complex mature root crowns, containing hundreds to thousands of roots. The orders of magnitude of increased complexity motivated unique solutions using both state-of-the-art techniques in computer graphics [38, 36, 37] and novel algorithms which eventually led to the development of TopoRoot. While we consider this first version as the foundation of several future planned advancements, discussed below, we were able to present here unprecedented fine-grained analysis of complex 3D root systems (on average 943 total roots [maximum of 2514]), 78 nodal roots (maximum of 126), 865 lateral roots (all classes combined; maximum of 2414) for the excavated maize roots, as computed by TopoRoot) that facilitates “apples to apples” comparisons with existing seedling phenotyping pipelines.

### Error analysis

The steps of our pipeline which are most prone to errors are topological simplification and skeletonization. The amount and quality of the topological repairs depends on the choices of the iso-values *k*, *s*, and *n*. If *k* and *n* are too close to *s*, then very little topological simplification will occur. On the other hand, if *k* and *n* are too far away from *s*, then such an aggressive setting may result in some topological features being removed in a geometrically suboptimal fashion. This is because we use a setting of [38] which uses the image intensity as guidance to determine where contents are added and removed from the iso-surface *s*, but these changes may not be smooth if the image values are not reliable. For example, cycles may be cut in the middle of branches, resulting in improper tracing of branches and potential double counting of the same root. Another step of our pipeline which is prone to errors is the cycle breaking portion of skeletonization. Decisions on where to break cycles rely on local angle continuity information near junctions, as well as the distance from the stem. However, if a root continues across many junctions where cycles pass through, then the root may accidentally be cut off early at one of these junctions. Our pipeline is sensitive to the amount of soil remaining in the sample, because soil often has a similar or higher intensity as the roots. If a large clump of soil is stuck onto the root, it may result in a thick region which is mistakenly identified as part of the stem. This problem can be avoided by thorough cleaning of samples prior to scanning, and by basic image thresholding prior to TopoRoot analysis. The final contrast will affect the ideal choices for *k* and *n* with respect to *s*; the greater the contrast, the smaller the gap between *k* and *n* need to be to incur the same amount of topological simplification.

The quality of the topological repairs may be improved by considering not only the image intensity when computing additions and removals of contents, but also incorporating geometric criteria such as the tubularity of a region to identify it as part of a root. This will allow for a greater amount of topological simplification without sacrificing geometric optimality. The suboptimal local decisions of cycle-breaking may be potentially improved by grouping cutting decisions together to produce a collective score. These groups can gradually be built from the bottom-up to produce a more globally optimal solution.

### Running time

On average, TopoRoot completes in 7 minutes and 13 seconds for each sample in the CT scan dataset (downsampled by a factor of 4 to the resolution of 369 x 369 x 465). Since this is much shorter than the time spent imaging and reconstructing one sample, TopoRoot is well suited for high-throughput analysis. The computation time is dominated by the first two steps, topological simplification (3 minutes and 6 seconds) and skeletonization (3 minutes and 44 seconds). The time complexity of both these steps may increase quickly with the image resolution. For example, running TopoRoot on the original CT volumes downsampled by a factor of 2 (instead of 4 as used in our validation), which results in volumes of resolution 737 x 737 x 931, would take 63 minutes and 39 seconds, with 32 minutes and 25 seconds spent on topological simplification and 29 minutes and 9 seconds on skeletonization. On the other hand, we have not observed a notable improvement in the accuracy of the nodal root count for this data set by reducing the downsampling factor from 4 to 2.

### Extensions

In addition to the per-level traits reported in this work, the hierarchy obtained by TopoRoots potentially enables computation of other fine-grained traits. For example, we are currently exploring the use of the hierarchy for computing whorls and the soil plane, which would in turn enable computation of traits such as inter-whorl distances, per-whorl measurements, and the numbers of nodal roots above and below the soil. Preliminary experiments show promising results of whorl detection by clustering the nodal roots along the stem. The soil plane can be potentially identified where a large cluster of 1st-order lateral roots appear along the direction of the stem.

While TopoRoot is designed for and validated on X-ray CT scans of excavated maize root crowns, the tool can be potentially adapted to other types of root systems and imaging modalities. For root crowns with multiple tillers (e.g., sorghum), we offer a mode of TopoRoot which extends the stem-detection heuristic (during the skeletonization step) by producing a stem path within each region of the skeleton above a given thickness threshold. Preliminary visual experiments show that TopoRoot’s multiple-tiller mode produces plausible hierarchies at a quality similar to that seen in the single-tiller mode (Figure 14). Further expanding the stem-detection heuristic to identify the primary root would make the pipeline applicable to taprooted systems as well. TopoRoot should work equally well for fine-grained analysis of 3D root system architecture reconstructions derived from in situ imaging methods such as X-ray CT, MRI, and optical imaging, provided they are first segmented from background (primarily soil and pot). Segmentation can be done using a variety of methods such as region-growing [1], tracking tubular features [26; 22], deep learning [29, 30], and semi-automatic annotation [3, 6]. Note that some of these methods produce a binary volume (e.g., region-growing) whereas others produce a probability density field (e.g., deep learning). Since TopoRoot requires a gray-scale intensity volume with three thresholds (shape, kernel and neighborhood), a binary segmentation will first need to be converted into a Euclidean distance field.

**Fig. 14.**
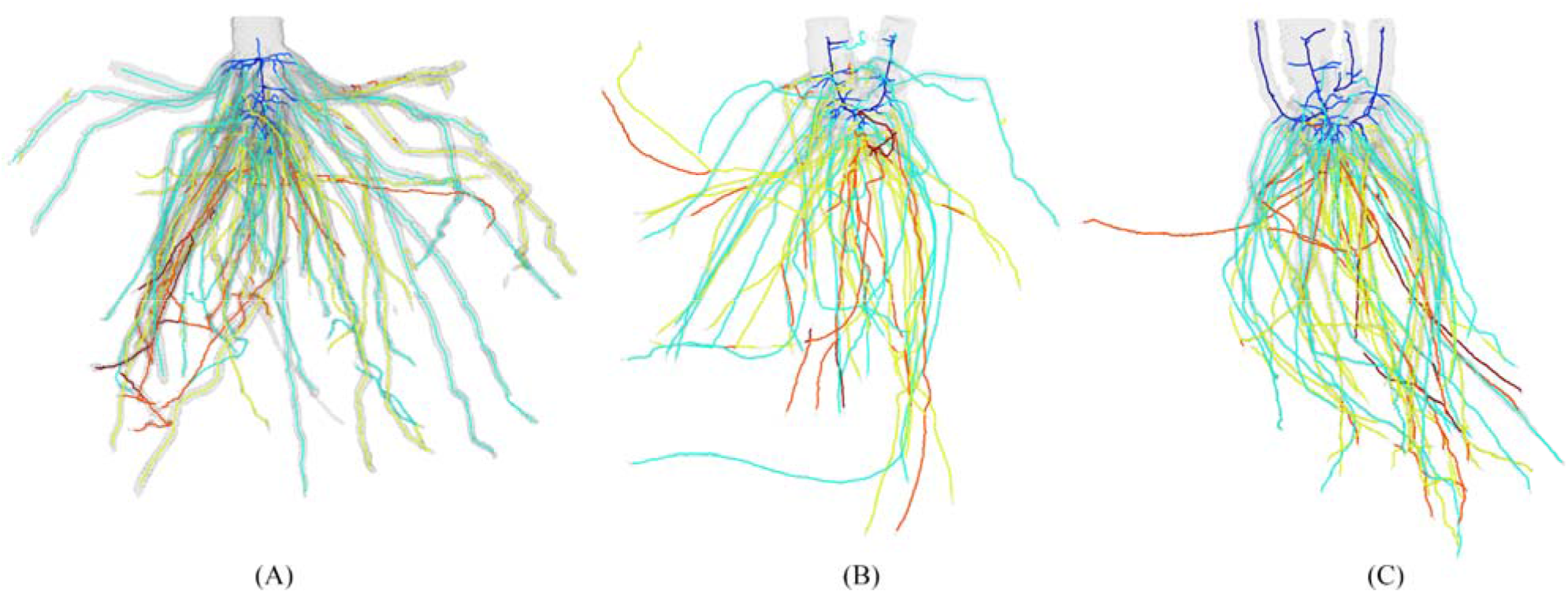
Hierarchies of sorghum roots computed by TopoRoot. The examples have (A) one tiller (B) two tillers and (C) four tillers. Hierarchy levels 0, 1, 2, 3 and 4 are colored dark blue, light blue, green, orange, and red.

### Software availability

TopoRoot is available for free at: https://github.com/danzeng8/TopoRoot Included in the page are instructions to run the software, and details on the formats of the input and output. Currently, the accepted inputs are either image slices (suffixed with .png) or .raw files, with a .dat accompanying the .raw file to specify the dimensions. TopoRoot is currently configured to build and run on Windows 10 machines, and has not been tested on other platforms. We plan on releasing a Linux build of TopoRoot in the future. A graphical user interface is also available on the page for visualizing the output hierarchy.

## Conclusions

We introduced TopoRoot, a high-throughput method for computing the root hierarchy and fine-grained root traits from a 3D image. TopoRoot specifically addresses topological errors, which are common in 3D imaging and segmentation and are barriers for obtaining accurate root hierarchies. Our method combines state-of-the-art methods developed in computer graphics with customized heuristics to compute a wide variety of traits at each level of the root hierarchy. When tested on both 3D scans of excavated maize root crowns and simulated root systems with artificially added noise, TopoRoot exhibits higher accuracy than existing tools (DynamicRoots and GiaRoots) in both coarse-grained and fine-grained traits. Furthermore, the efficiency and automation of TopoRoot makes it ideal for a high-throughput analysis pipeline, and the results are readily compatible with the Root System Markup Language (RSML;[13]), and major plant structural-functional modelling frameworks such as CRootBox [25] and OpenSimRoot [24].

## List of Abbreviations

RSA: Root system architecture
CT: Computed Tomography

## Declarations

### Ethics approval and consent to participate

Not applicable

### Consent for publication

Not applicable

### Availability of data and materials

The datasets generated and analysed during the current study are available in the TopoRoot Github repository: https://github.com/danzeng8/TopoRoot

### Authors’ Contributions

DZ was the primary developer and coder of TopoRoot. DZ wrote the manuscript with contributions from TJ, CNT, NJ, ML, EC, DL, YJ, and HS. DZ, TJ, and CNT designed the study and experiments. DZ performed experiments and analyzed the data. YJ developed the graphical user interface with contributions from DZ. All authors have read and approved the final manuscript.

## Acknowledgements

We would like to thank Tiffany Hopkins, Dhineshkumar Thiruppathi, Elisa Morales, Mitchell Sellers, Shayla Gunn, August (Gus) Thies, Keith Duncan, and Tim Parker (Donald Danforth Plant Science Center) for the planting, harvesting, imaging, and collection of hand measurements on maize roots, and Gustavo Gratacos (Washington University) and Yajie Yan (Facebook) for their valuable insights on the cycle breaking algorithm and geometric skeleton computation. We thank Mon-Ray Shao (Donald Danforth Plant Science Center) for the discussion on future applications to sorghum and other species.

## Authors’ information

Affiliations

**Department of Computer Science and Engineering, Washington University in St. Louis, St. Louis, USA**

Dan Zeng, Tao Ju, Yiwen Ju

**Donald Danforth Plant Science Center, St. Louis, USA**

Christopher Topp, Mao Li, Ni Jiang

**Department of Computer Science, Saint Louis University, St. Louis, USA**

Erin Chambers, David Letscher, Hannah Schreiber

## Supplementary Information

**Supplementary Table 1:** TopoRoot’s computed traits.

| Groups | Trait | Description |
| --- | --- | --- |
| Global traits | Total length | Sum of the lengths of stem, nodal roots, and lateral roots. |
|  | Number of roots | Total number of roots across all hierarchy levels. |
|  | Average length | Total length divided by number of roots. |
| Stem traits | Average stem thickness | Average thickness of the vertices along the stem path. |
|  | Stem length | Length of the stem path. |
| Per-level traits | Level $n$ root count | The number of level $n$ roots. |
| | Total level $n$ root length | Sum of the lengths of level $n$ roots. The length is the skeleton distance from the beginning of the root to its tip. |
| | Average level $n$ root length | Total level $n$ root length divided by level $n$ root count. |
| | Level $n$ root tortuosity | Length of a root (skeleton distance) divided by the Euclidean distance from the beginning to the tip, averaged across all level $n$ roots. |
| | Level $n$ root thickness | Thickness associated with the skeleton vertices in the root, averaged across all level $n$ roots. |
| | Number of level $n$ root children | Number of level 2 roots divided by number of level 1 roots. |
| | Level $n$ root tip angle | Angle between the stem direction and the vector from the beginning to the tip of a root, averaged across all level $n$ roots. |
| | Level $n$ root emergence angle | Angle between the stem direction and the vector from the beginning to 30 vertices along the skeleton of a root, averaged across all level $n$ roots. |
| | Level $n$ root midpoint angle | Angle between the stem direction and the vector from the beginning of a root to the halfway point of the root, averaged across all level $n$ roots. |
| Aggregated lateral root traits | Total lateral length | Sum of the lengths of lateral roots whose hierarchy level is greater than or equal to 2. |
|  | Number of lateral roots | Number of lateral roots whose hierarchy level is greater than or equal to 2. |
|  | Average lateral root length | Average length of lateral roots whose hierarchy level is greater than or equal to 2. |

**Supplementary Fig. S1.**
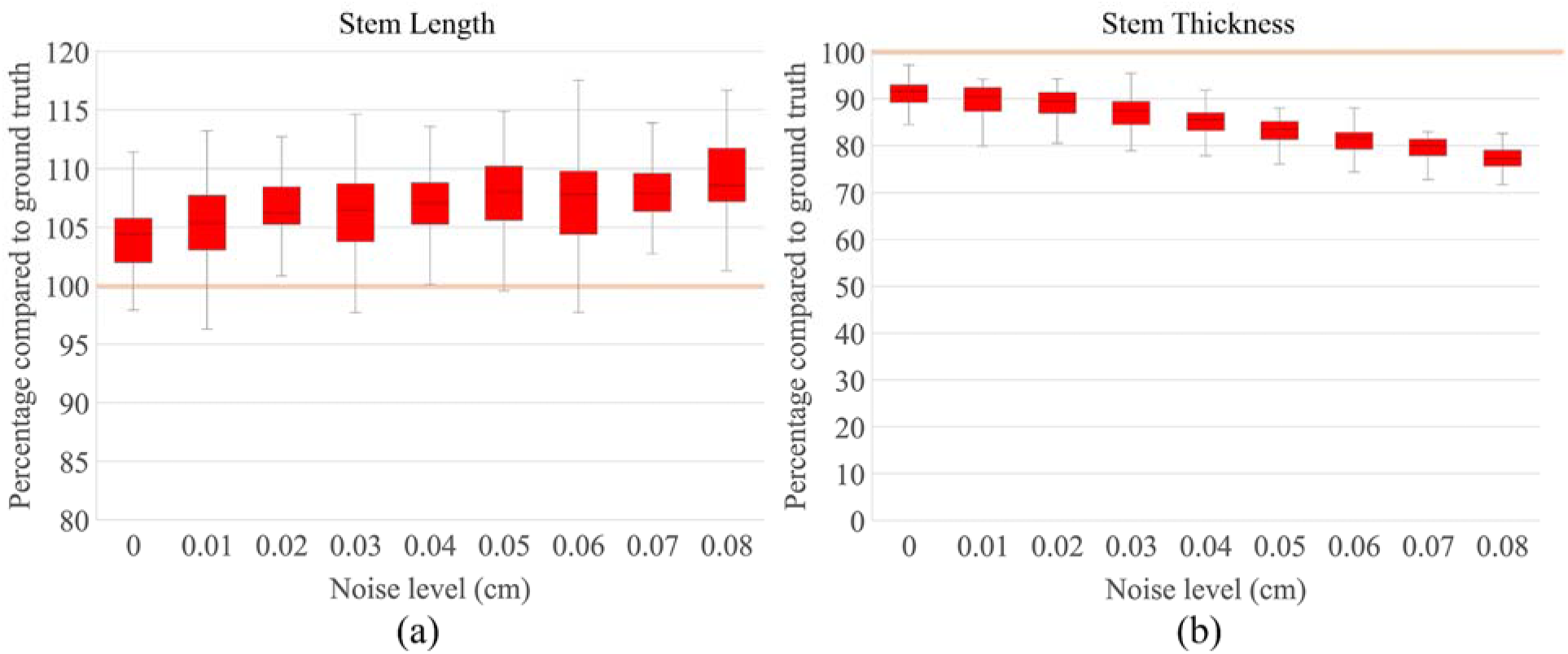
Evaluating the accuracy of computing stem traits by TopoRoot. The tan line represents the ground truth, and the result of TopoRoot is computed as a percentage of the ground truth.

**Supplementary Fig. S2.**
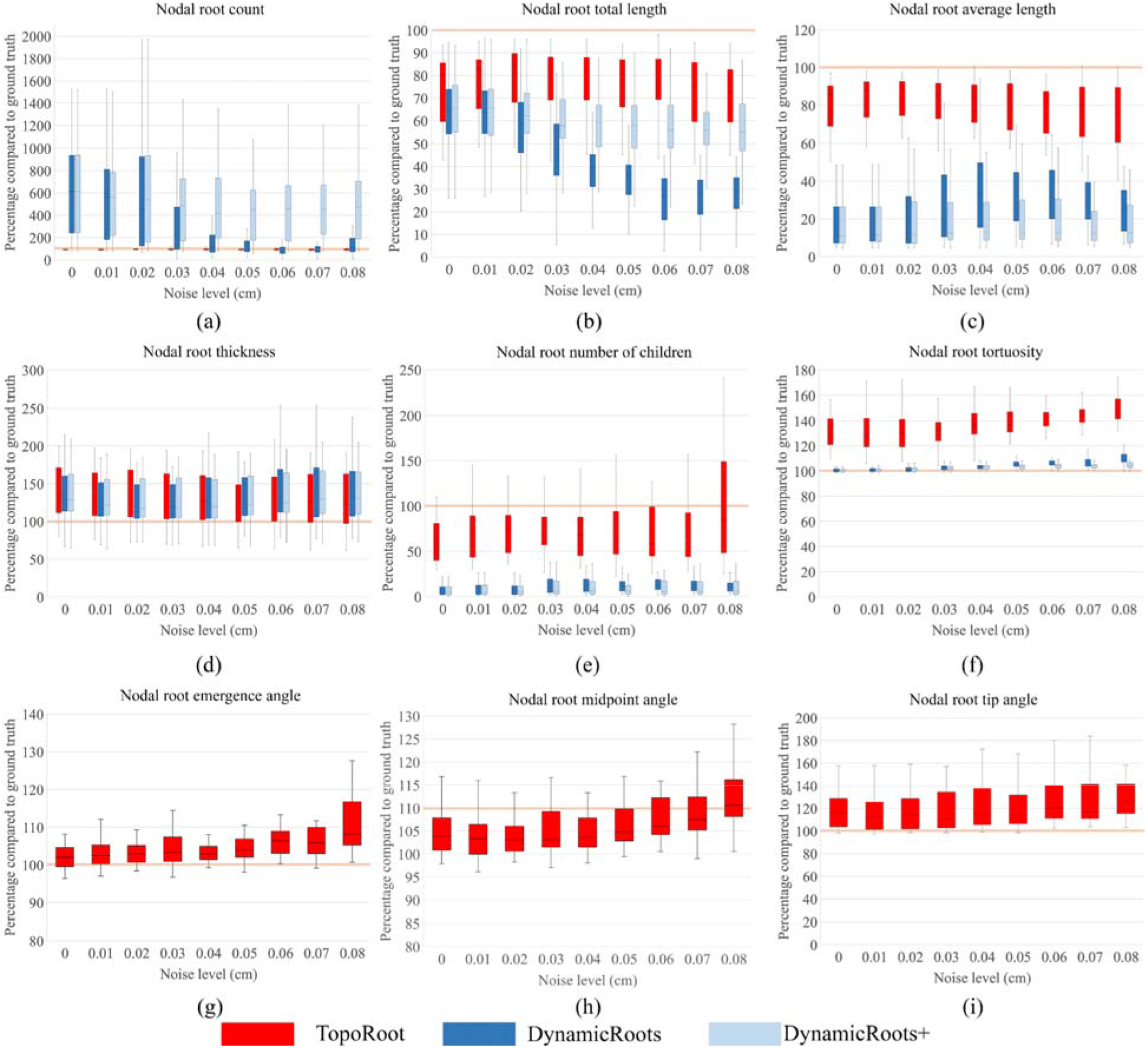
Comparing the accuracy of computing nodal root traits between TopoRoot, DynamicRoots, and DynamicRoots+ across 55 models. The tan line represents the ground truth, and the results of all methods are computed as a percentage of the ground truth.

**Supplementary Fig. S3.**
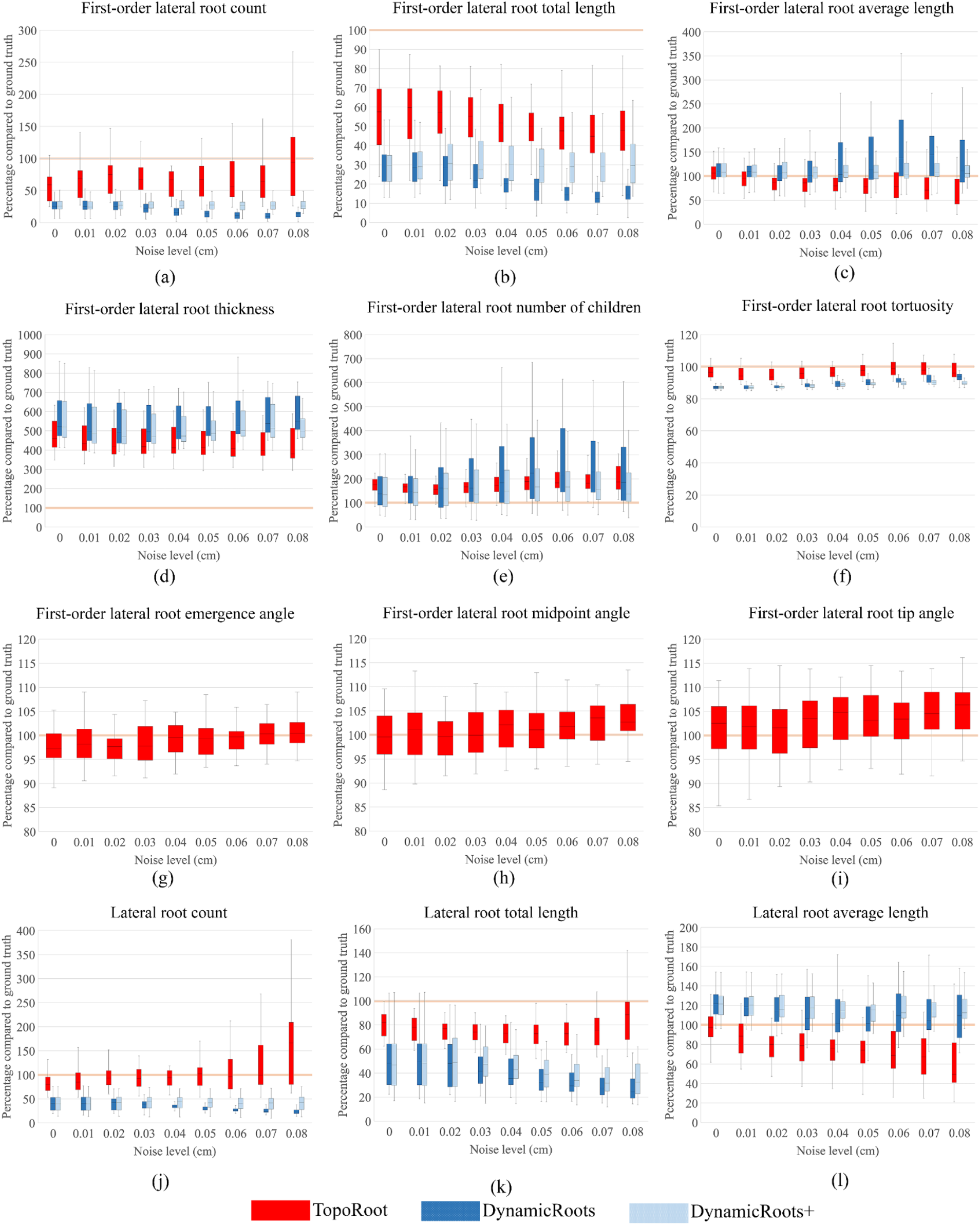
Comparing the accuracy of computing lateral root traits between TopoRoot, DynamicRoots, and DynamicRoots+ across 55 models. The tan line represents the ground truth, and the results of all methods are computed as a percentage of the ground truth.

**Supplementary Fig. S4.**
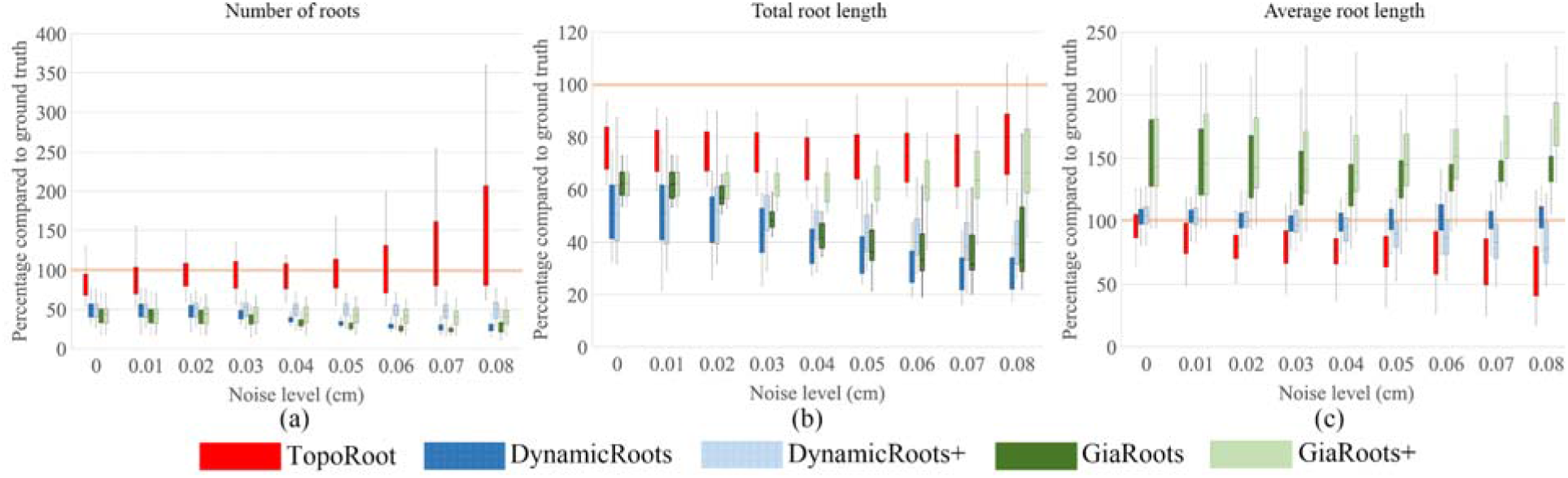
Comparing the accuracy of computing global traits between TopoRoot, DynamicRoots, DynamicRoots+, GiaRoots, and GiaRoots+ across 55 models.The tan line represents the ground truth, and the results of all methods are computed as a percentage of the ground truth.

