## Supplementary figures and images for "TopoRoot: A method for computing hierarchy and fine-grained traits of maize roots from X-ray CT images"

### Supplementary Figure 1

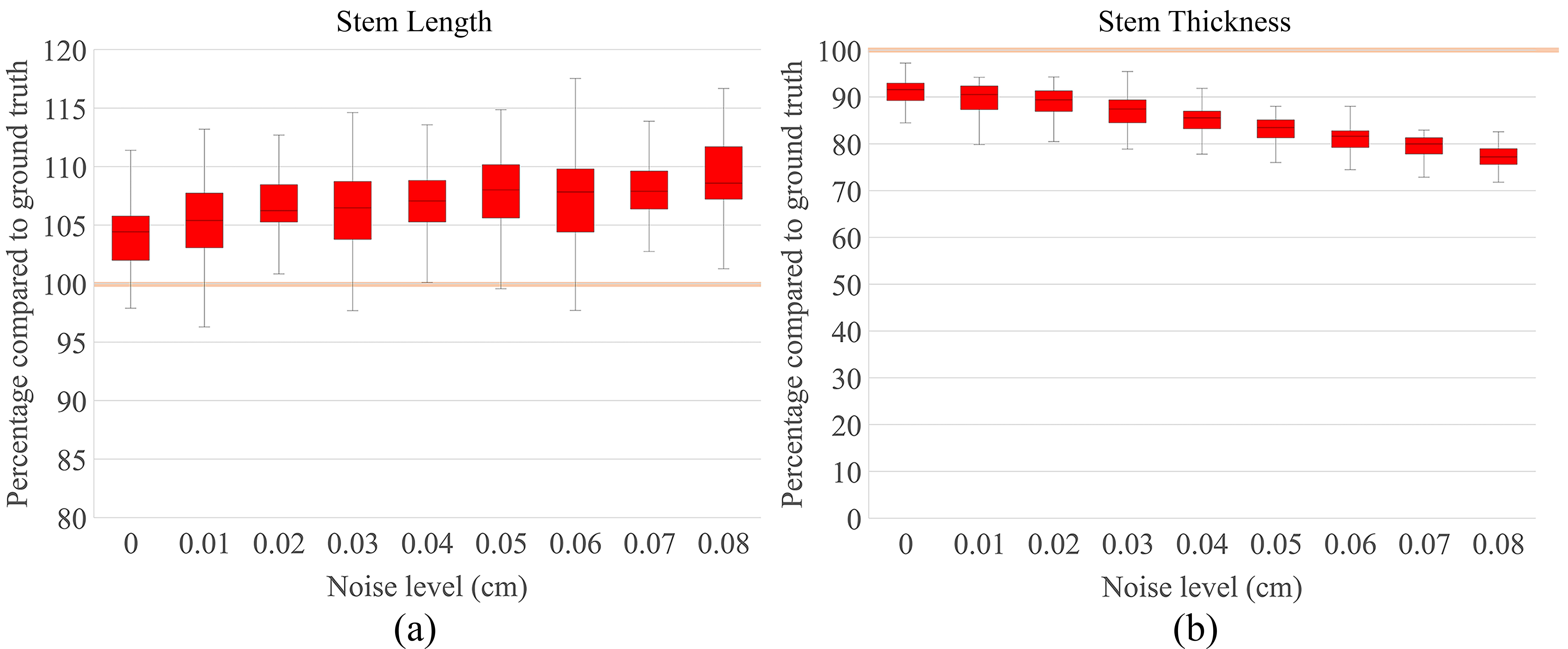

### Supplementary Figure 2

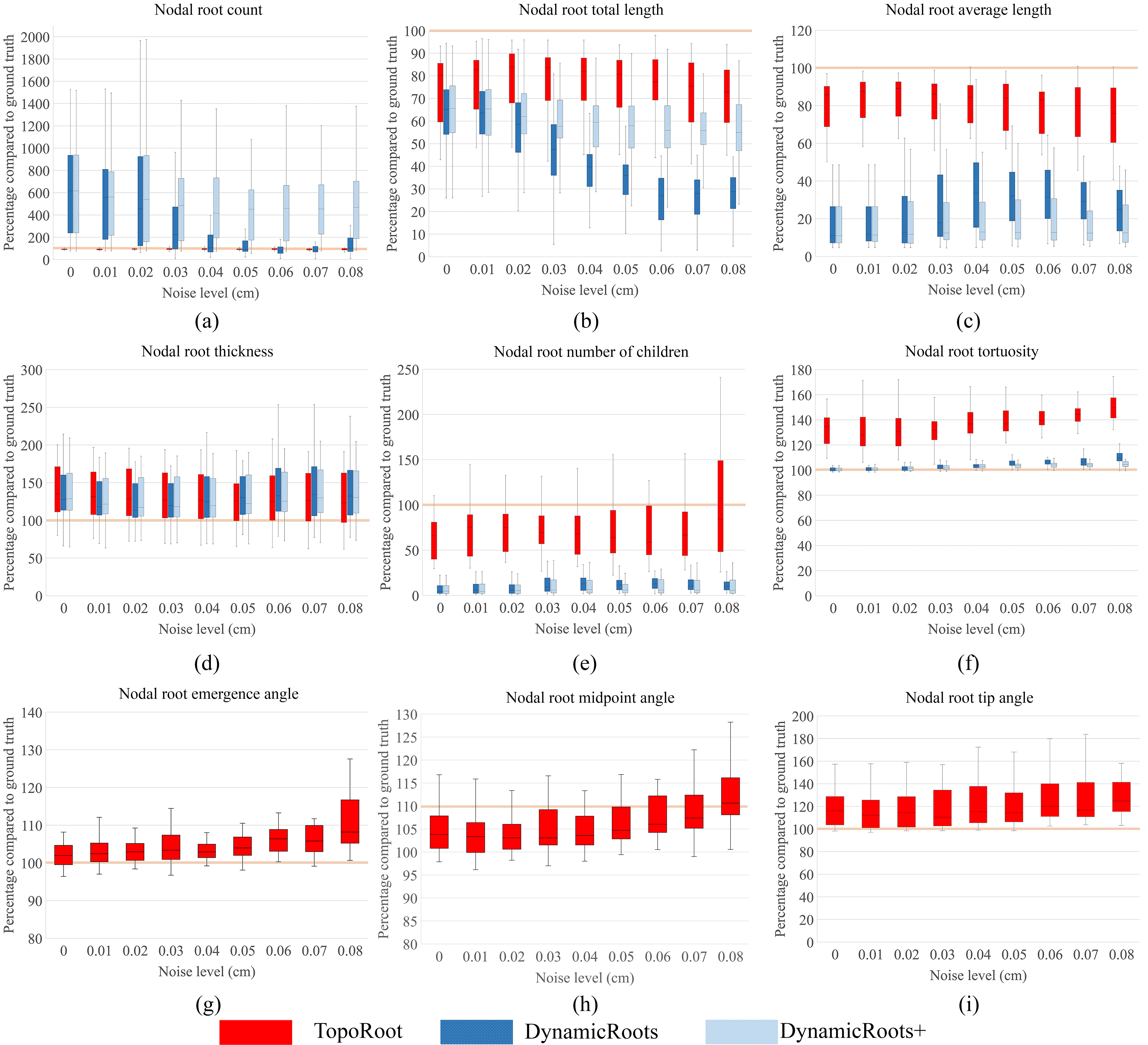

### Supplementary Figure 3

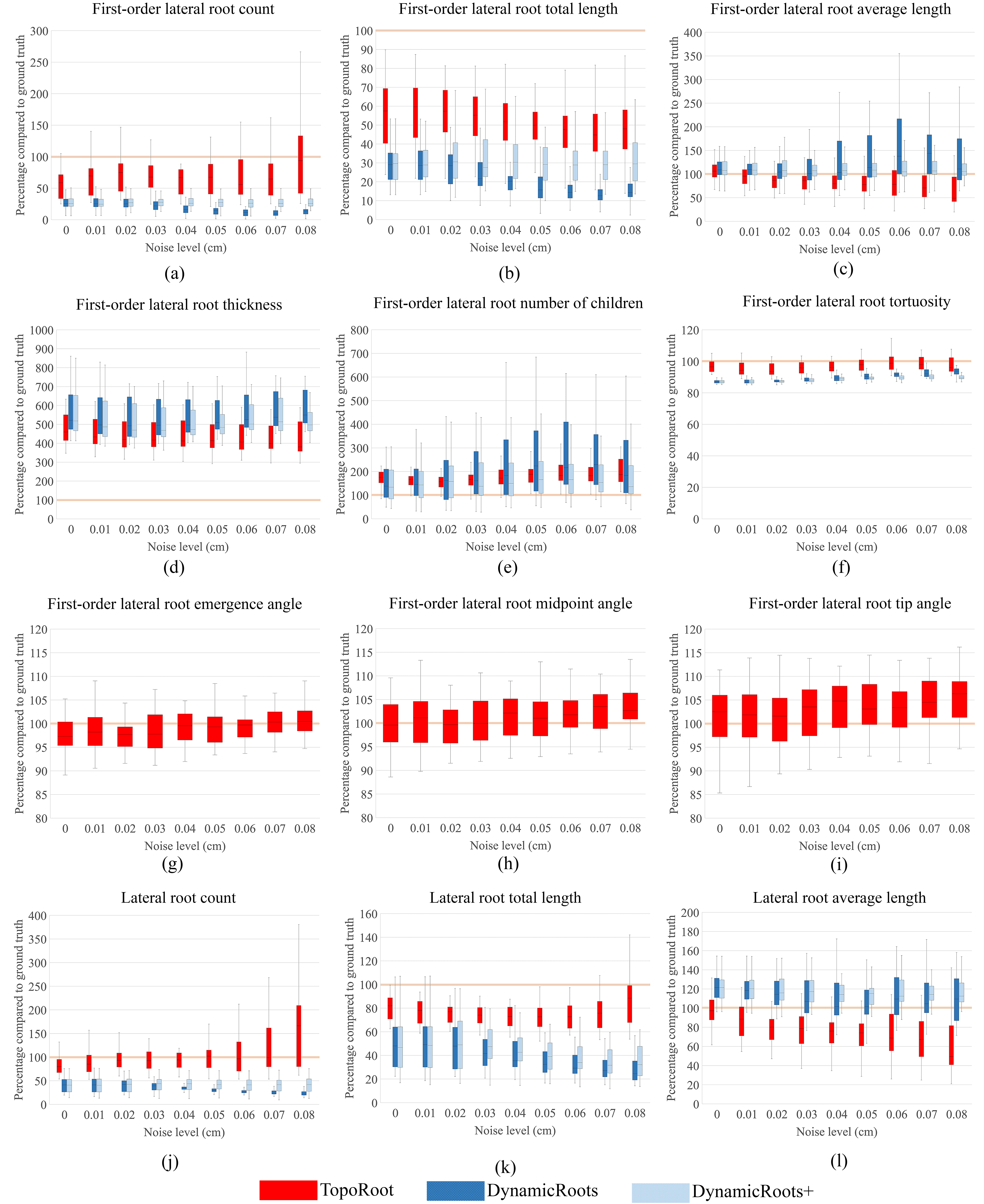

### Supplementary Figure 4

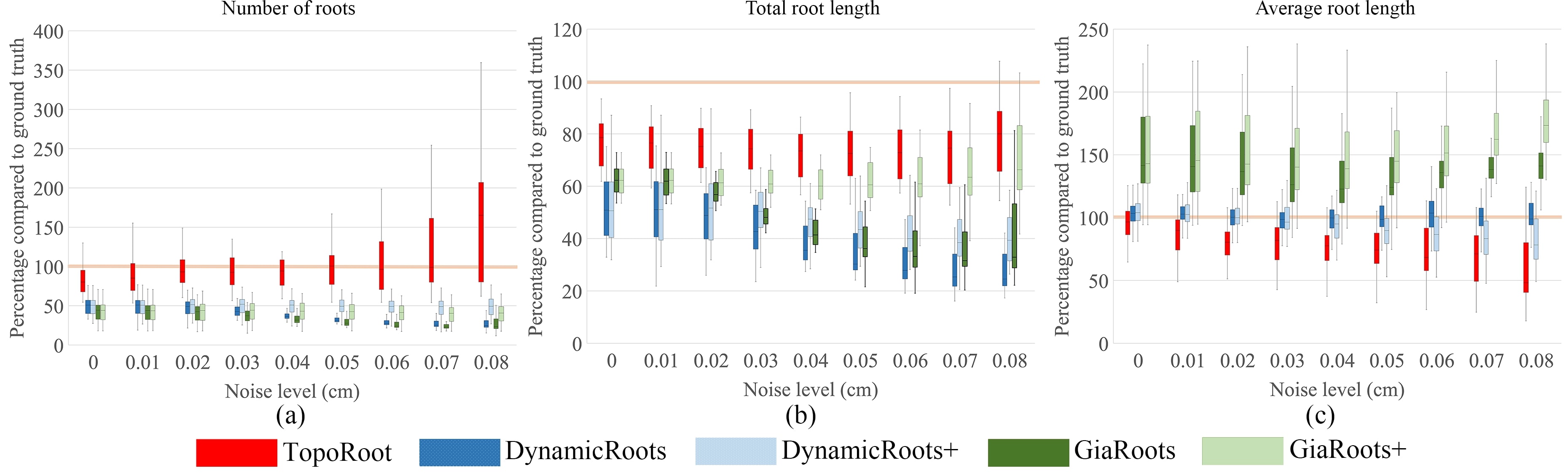
